# Anatomy of the mandibular corpus of extant cercopithecids : taxonomy and variation

**DOI:** 10.1101/2024.03.31.587458

**Authors:** Laurent Pallas, Masato Nakatsukasa, Yutaka Kunimatsu

## Abstract

This study aims to discriminate cercopithecid taxa of higher taxonomic levels (subfamily, tribe, subtribe, and genus) on the basis of corpus shape in transverse cross-section at the M_1_-M_2_ junction and to assess its variation using 2D geometric morphometrics. Specifically, we evaluated the effect of allometry and sexual dimorphism on differences in corpus shape at interspecific and intraspecific levels, respectively. We also investigated whether corpus variation among cercopithecids was following Brownian motion using Pagel’s λ. Taxonomic discrimination and sexual dimorphism were established using Analysis of Variance on Principal Component scores. Allometry was studied using phylogenetic least-squares regressions and partial least-squares regressions. We demonstrated that, using corpus shape, extant cercopithecids can be significantly discriminated at the subfamilial, tribal, and subtribal levels. In addition, the main axis of variation of the Principal Component Analysis follows a distribution expected under Brownian motion, validating the presence of a phylogenetic signal in corpus shape. Colobines exhibit a robust corpus (superoinferiorly short and transversely broad) with large lateral prominences while cercopithecines have a gracile corpus (superoinferiorly long and transversely thin in its distal portion) with marked corpus fossae in African papionins. Exception to the typical subfamilial or tribal shape pattern exist, with the best examples being *Trachypithecus*, *Presbytis* and *Pygathrix* within colobines, *Allenopithecus* within Cercopithecini, and *Macaca*, *Theropithecus* and *Cercocebus* within Papionini. Sexual dimorphism is a confounding factor in shape discrimination, as there are significant differences between sexes, notably in *Papio anubis*, *Nasalis larvatus* and *Procolobus verus* . Intriguingly, sexual dimorphism in corpus shape does not seem to follow the dimorphism deduced in canine and molar crown dimensions. This discrepancy is illustrated by the low degree of dimorphism in corpus shape in *Piliocolobus badius*, despite dimorphic canine and molar dimensions. Overall, our findings concerning corpus shape variation in cercopithecids will greatly benefit to paleontological studies that seek to identify taxa in the fossil record, and to neontological studies aiming to explore the ecomorphological value of the cercopithecid mandible.

## INTRODUCTION

The shape of the extant catarrhines corpus is a useful taxonomic discriminator and has been implicated in taxonomic discrimination among extant cercopithecoids (Takahashi and Pan, 1994; Jablonski et al., 1998; Daegling and McGraw, 2001; Pan et al., 2002, 2008; Wright et al., 2008) and hominoids (Daegling, 1989; Humphrey et al., 1999; Taylor, 2002; Taylor and Groves, 2003; Taylor, 2006a; Pitirri and Begun, 2019; Pitirri et al., 2020). The sources of variation in corpus shape, notably sexual dimorphism and ontogeny, were clarified in extant hominoids (Wood et al., 1991; Daegling, 1996; Brown, 1997; Taylor, 2006b; Singh, 2014; Pitirri and Begun, 2020). Such data are of paramount value to interpret the mandibular hominoid fossil record (White and Johanson, 1982; Chamberlain and Wood, 1985; Lockwood et al., 1996; White et al., 2000; Fabbri, 2006; Skinner et al., 2006; Lague et al., 2008; Haile-Selassie et al., 2015, 2022; Ioannidou et al., 2022). Despite a rich fossil record (Freedman, 1957; Delson, 1973; Leakey, 1982; Benefit and Pickford, 1986; De Bonis et al., 1990; Frost and Delson, 2002; Leakey et al., 2003; Hlusko, 2006, 2007; Jablonski and Leakey, 2008; Jablonski et al., 2008a, 2008b; Nakatsukasa et al., 2010; Pallas, 2019; Gommery et al., 2022) and the great taxonomic and functional diversity of its extant representatives (Groves and Kingdon, 2013; Rowe and Jacobs, 2016a), a thorough examination of the corpus shape of extant Old World Monkeys (Cercopithecidae) on a large taxonomic scale is lacking, preventing the extrapolation of any taxonomical and ecomorphological considerations in extant and fossil representatives of this group.

Approximately 23 genera of cercopithecids are documented in Africa and Eurasia (Groves and Kingdon, 2013a), with diet ranging from specialized follivory to omnivory. Cercopithecids can be divided into two subfamilies, Colobinae (leaf-eating monkeys) and Cercopithecinae (cheek-pouch monkeys). This taxonomic differentiation is followed by distinct adaptive strategies, with a rather ecclectic and versatile diet in cercopithecines and a more specialized diet oriented towards follivory and granivory in colobines (Kingdon and Groves, 2013a; Kingdon and Groves, 2013b; Rowe and Jacobs, 2016a; Rowe and Jacobs, 2016b; Rowe and Jacobs, 2016c). Cercopithecids are also used as model organisms to understand the complex interplay observed between diet, dietary behaviors, mastication biomechanic and mandibular anatomy (Hylander, 1979; Bouvier, 1986; Daegling, 1993, 2002; Ravosa, 1996; Jablonski et al., 1998; Vinyard and Ravosa, 1998; Daegling and McGraw, 2001, 2007; Daegling et al., 2009; Panagiotopoulou and Cobb, 2011; McGraw and Daegling, 2020; Panagiotopoulou et al., 2020). Previous analysis of the shape of the cercopithecid corpus used raw estimates of shape in the form of width and height (Hylander, 1979; Smith, 1983; Bouvier, 1986; Takahashi and Pan, 1994; Ravosa, 1996; Jablonski et al., 1998; Pan et al., 2002, 2008) or biomechanical parameters such as corpus cross-sectional area (Smith, 1983; Daegling and McGraw, 2001; Daegling, 2007). Although useful to discriminate extant and fossil cercopithecid taxa, these parameters, usually in the form of a ratio also referred as mandibular robusticity (e.g., Pallas et al., 2019), do not fully characterize the complex shape of the cercopithecid corpus, including the development of the lateral prominence, submandibular fossae, and corpus fossae, which are considered taxonomically diagnostic. Indeed, corpus anatomy has been used in diagnosis of fossil (Freedman, 1957; Leakey et al., 1983; Benefit and Pickford, 1986; Frost and Delson, 2002; Leakey et al., 2003; Hlusko, 2006,2007; Pallas et al., 2019; Gommery et al., 2022) and extant cercopithecids (Frost, 2001; Groves, 2007; Gilbert et al., 2018) but the level of variation, and particularly the influence of sexual dimorphism, in traits such as the lateral prominences, for example, is unknown. Without such data, taxonomic discrimination in the fossil record is a difficult task.

Here, we used 2D morphometric analysis (landmarks and sliding semilandmarks) to quantify the corpus shape of 22 genera of cercopithecids at the M_1_-M_2_ level. Principal Component Analysis (PCA) was employed to visualize and order shape variation, and significant differences in Principal Component scores between taxa were investigated using Analysis of Variances (ANOVAs). We assessed at the tribal and subtribal levels, differences in allometric trajectories of corpus shape using centroid size as a body mass proxy. Partial least squares regressions (PLS) were used to model allometry and significant differences in slope values between taxa were then investigated. In addition, we also computed phylogenetic generalized least squares (PGLS) regressions to account for the confounding effect of phylogeny on allometry. As one of the main sources of shape variation, we tested for the effect of sexual dimorphism by assessing significant differences between male and female specimens of a set of cercopithecid species known to differ in their magnitude of sexual dimorphism. Finally, we evaluated whether the distribution of corpus shape data is structured by phylogeny as expected under Brownian motion using Pagel’s λ. Our results are a first step towards a better understanding of the phenetic diversity of extant cercopithecids, and subsequently pave the way for finer ecomorphological analyses on extant cercopithecids, as well as a reassessment of the taxonomy of fossil cercopithecids.

## MATERIAL AND METHODS

### NEONTOLOGICAL SAMPLE

We collected data on 795 specimens from 72 species of 22 cercopithecid genera : Allenopithecus (*n* = 16), Allochrocebus (*n* = 4), Cercocebus (*n* = 16), Cercopithecus (*n* = 64), Chlorocebus (*n* = 24), Colobus (*n* = 103), Erythrocebus (*n* = 10), Lophocebus (*n* = 25), Macaca (*n* = 188), Mandrillus (*n* = 7), Miopithecus (*n* = 3), Nasalis (*n* = 48), Papio (*n* = 69), Piliocolobus (*n* = 42), Presbytis (*n* = 70), Procolobus (*n* = 25), Pygathrix (*n* = 5), Rhinopithecus (*n* = 2), Semnopithecus (*n* = 15), Simias (*n* = 25), Theropithecus (*n* = 15), and Trachypithecus (*n* = 18). Detailed information on sexes, accession numbers, taxonomy of the specimens as well as the provenance of the 3D models is provided in SOM Table S1.

### MORPHOMETRIC PROTOCOL

We obtained photographs of the coronal cross-sections set at the M_1_/M_2_ junction using Avizo v.7.0 (Thermo Fisher Scientific, Waltham). First, the mandible was aligned in lateral view by setting the alveolar process parallel to the transverse plane using the trackball function of Avizo. Second, a cross-section was set at M_1_/M_2_ junction. Third, the mandible was rotated using the trackball function of the software so that the genioglossal fossa was aligned with the sagittal plane and fourth, a photograph of the cross-section of the corpus was then taken.

We digitized *n* = 3 landmarks and *n* = 75 sliding semi-landmarks using tpsDig2 v.2.32. The outline of the corpus was digitized using semilandmarks and we used the “Resample curve” function of tpsDig2 to distribute the semilandmarks at an equidistant length along the outline of the corpus. The position of the landmarks and semi-landmarks is illustrated in Figure 1. The three landmarks represent the superior (both lingual and labial) and inferior aspect of the corpus. Wherever possible, the right corpus has been digitized. In cases where the right corpus was damaged, we flipped the left corpus horizontally and digitized it has a right corpus.

**Figure 1:**
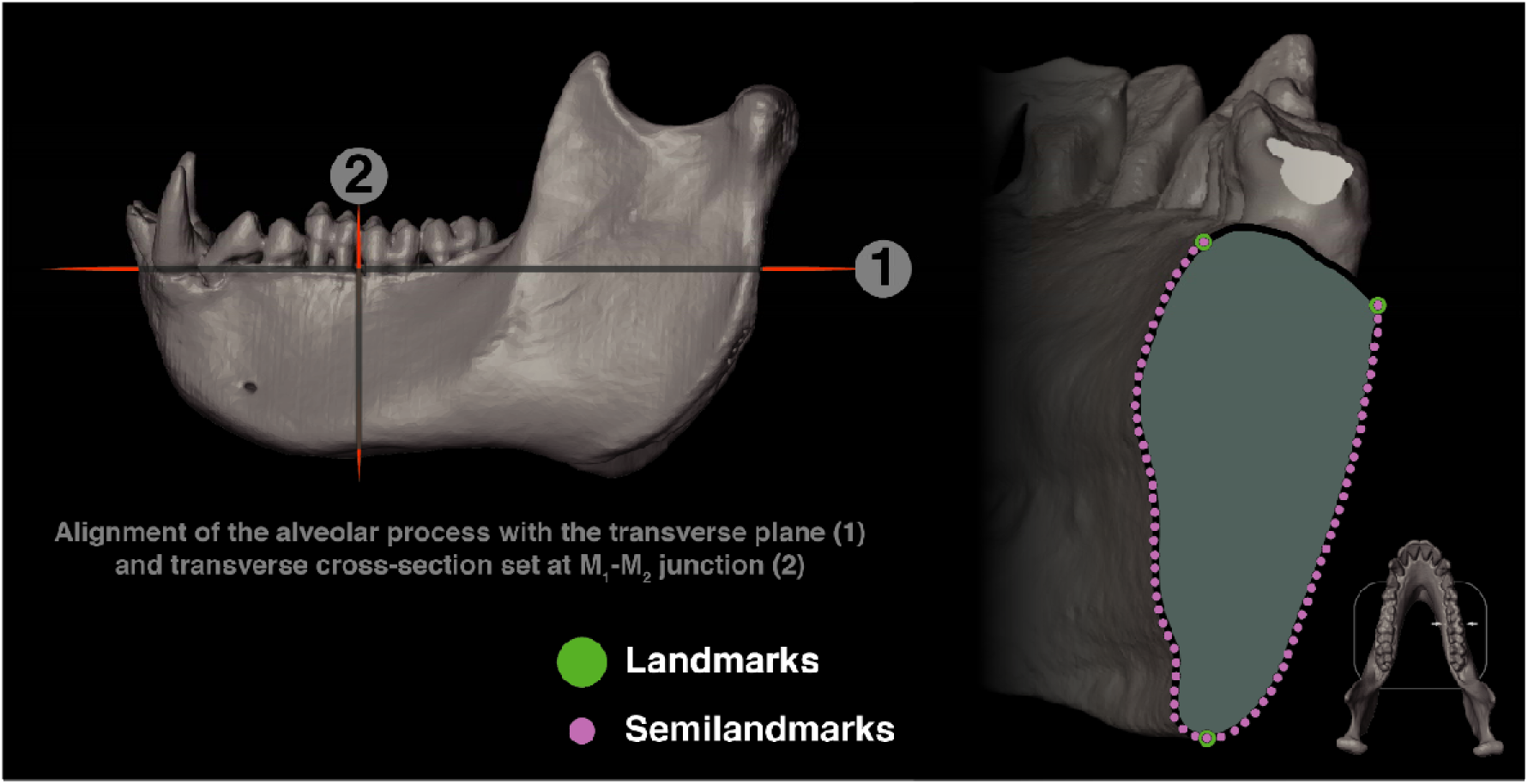
Illustration of the acquisition protocol of corpus cross-sections at the M _1_-M_2_ junction and of landmarks and semi-landmarks placement.

### STATISTICAL ANALYSES

All statistical analyses were performed using R v. 3.5.0 (R Core Team, 2021). Prior to the statistical tests, we tested the normal distribution and homoscedasticity of the model residuals using Shapiro-Wilk and Bartlett’s test (SOM Table S1 and S2). We used non-parametrical tests when the null-hypothesis of homogeneity of the variances (i.e., homoscedasticity) was not verified. We set the level of significant of the statistical tests at 5% and 1% (significant differences) as well as 0.1% (highly significant differences).

#### Generalized Procrustes Analysis and shape coordinates

We imported landmarks and semidlandmarks 2D coordinates in R. We defined semilandmarks using the define.sliders() function of the ‘geomorph’ package, and then subjected the raw coordinates to a Procrustes analysis using the gpagen() function of ‘geomorph’. In order to visualize the mean shape of selected taxa, we used the mshape() function of ‘geomorph’. We computed mean shape per genera only for genera that are represented by more than 5 specimens in our dataset (SOM Table S1).

#### Principal Component Analysis

We explored and ordered variation in corpus shape using a Principal Component Analysis (PCA) on the Procrustes shape variables obtained using the gpagen() function. The PCA was computed using the gm.prcomp() function of ‘geomorph’.

#### Significant differences in multivariate data

As the primary objective of this study is to discriminate specimens based on taxonomy and sexes, we used Analysis of Variance (ANOVA), along with Tukey’s Honest Significant Difference (HSD) post-hoc test, to identify significantly distinct pairs of taxa and significant differences between male and female specimens among given taxa based on Principal Component (PC) scores. When homoscedasticity was not verified, we used the Kruskall- Wallis test, combined with a Dunn’s post-hoc test. Parameters of the ANOVA’s and Tukey’s HSD models were calculated using the aov() and Tukey(HSD) functions of the ‘stats’ package. The kruskal.test() and dunnTest() functions of the ‘stats’ and ‘FSA’ packages were used to compute the parameters of the Kruskall-Wallis’s and Dunn’s tests.

The confounding effect of sexual dimorphism (i.e., significant differences between male and female specimens of a given taxa) has been evaluated on a limited number of species. We used Gilbert and Grine (2009) criteria of about 15 specimens to select such species. Therefore, sexual dimorphism is evaluated in the following taxa: *Colobus polykomos*, *Colobus guereza*, *Piliocolobus badius*, *Procolobus verus*, *Presbytis bicolor*, *Nasalis larvatus*, *Cercopithecus mitis*, *Chlorocebus aethiops*, *Macaca fascicularis*, *Macaca fuscata*, and *Papio anubis* . This includes taxa with marked sexual dimorphism in body mass, dental dimensions, and symphyseal shape (e.g., *P. anubi*s and *N. larvatus*) and poorly dimorphic species (e.g., *Pre. bicolor*).

#### Comparative phylogenetic analyses

We evaluated the null-hypothesis that our dataset is structured by phylogeny assuming a Brownian Motion (BM) model. In this framework, the difference in PCA scores of cercopithecid taxa is proportional (supposing that the generation times are homogeneous) to their divergence times. To test this hypothesis, we evaluated the effect of phylogeny on the structuration of our dataset using Pagel’s λ, calculated by maximum likelihood using the phylosig() function of the ‘phytools’ package. We assembled a phylogenetic tree using data (i.e., branch lengths, divergences dates and topologies) from Perelman et al. (2011), with minor modifications of branch lengths and divergence dates according to Ting (2008) and Liedigk et al. (2012) data. Compared to the tree of Perelmann et al. (2011), we added *Simias concolor* as a sister taxon to *Nasalis larvatus* after the divergence date from Liedigk et al. (2012) and phylogenetic hypotheses of Whittaker et al. (2006) and Liedigk et al. (2012). We used Ting (2008) divergence date for the *Colobus* – *Piliocolobus*/*Procolobus* dichotomy.

We used the available R script from Barr (2020) for the phylomorphospace shown in Figure 2.

**Figure 2:**
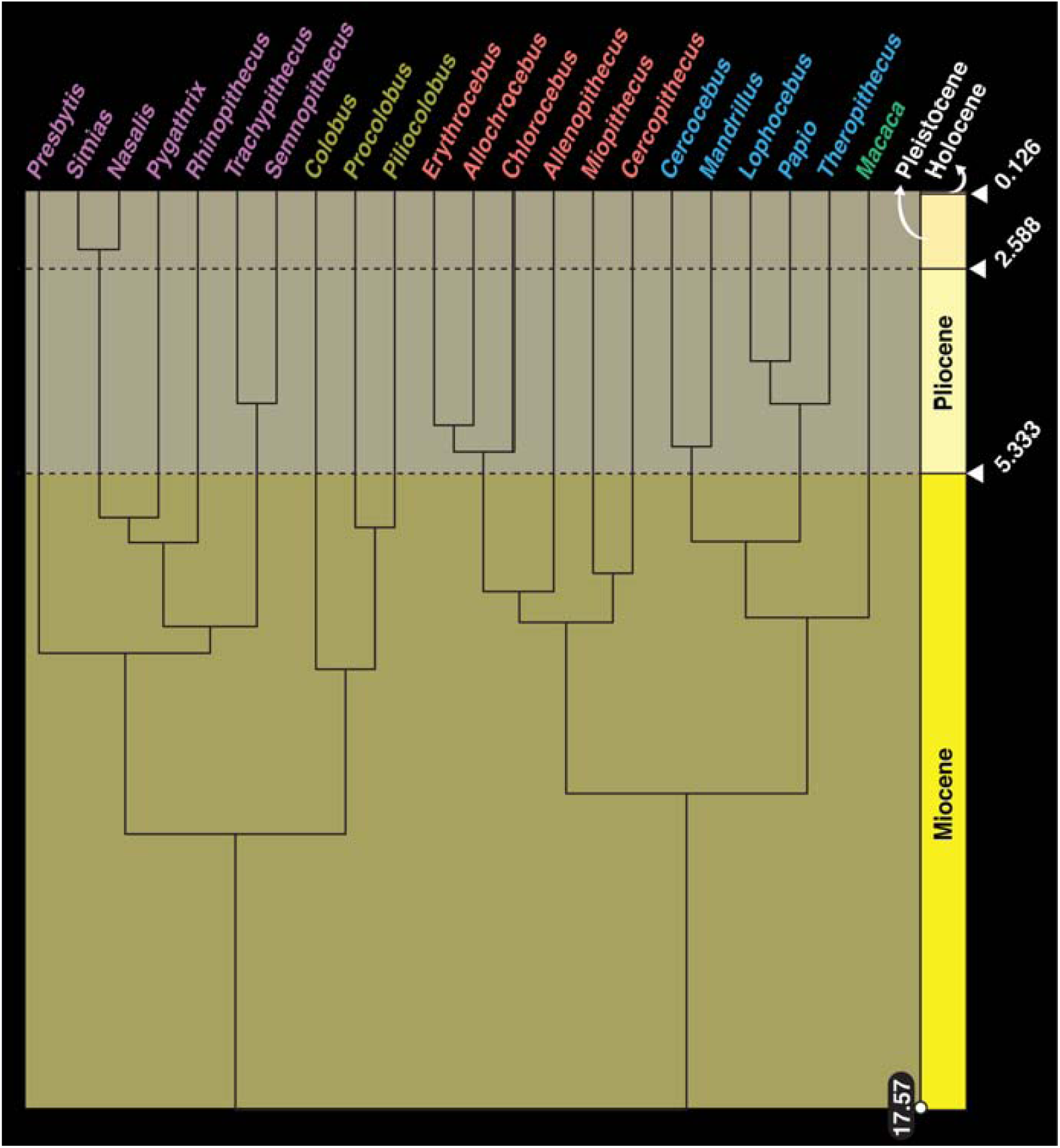
Chronogram used in the assessment of the phylogenetic signal embedded in PC scores.

#### Allometry

We evaluated the effect of size in shape differences between taxa (interspecific allometry) using both phylogenetic least squares regressions (PGLS) and partial least squares (PLS) regressions. We modeled the size using the centroid size obtained from each specimen after running the Procrustes analysis. Allometry was hence modeled as the natural logarithm of the PC scores as the response variable and the natural logarithm of the centroid size as the predictor variable. An arbitrary value of one was added to each PC scores to log transform them. PGLS regressions were established using the pgls() function of the ’caper’ package with λ transformation computed using maximum likelihood. PLS regressions were calculated with the lm() of the ’stats’ package.

## RESULTS

### PCA SCORES AND OVERALL MORPHOMETRIC DIFFERENCES

PC1 and PC2 combined account for nearly 66.0% of the variance (Figure 3). Cercopithecines and colobines can differentiated on PC1 (ca. 44.0% of the variance), with colobines presenting negative scores and cercopithecines positive scores.

**Figure 3:**
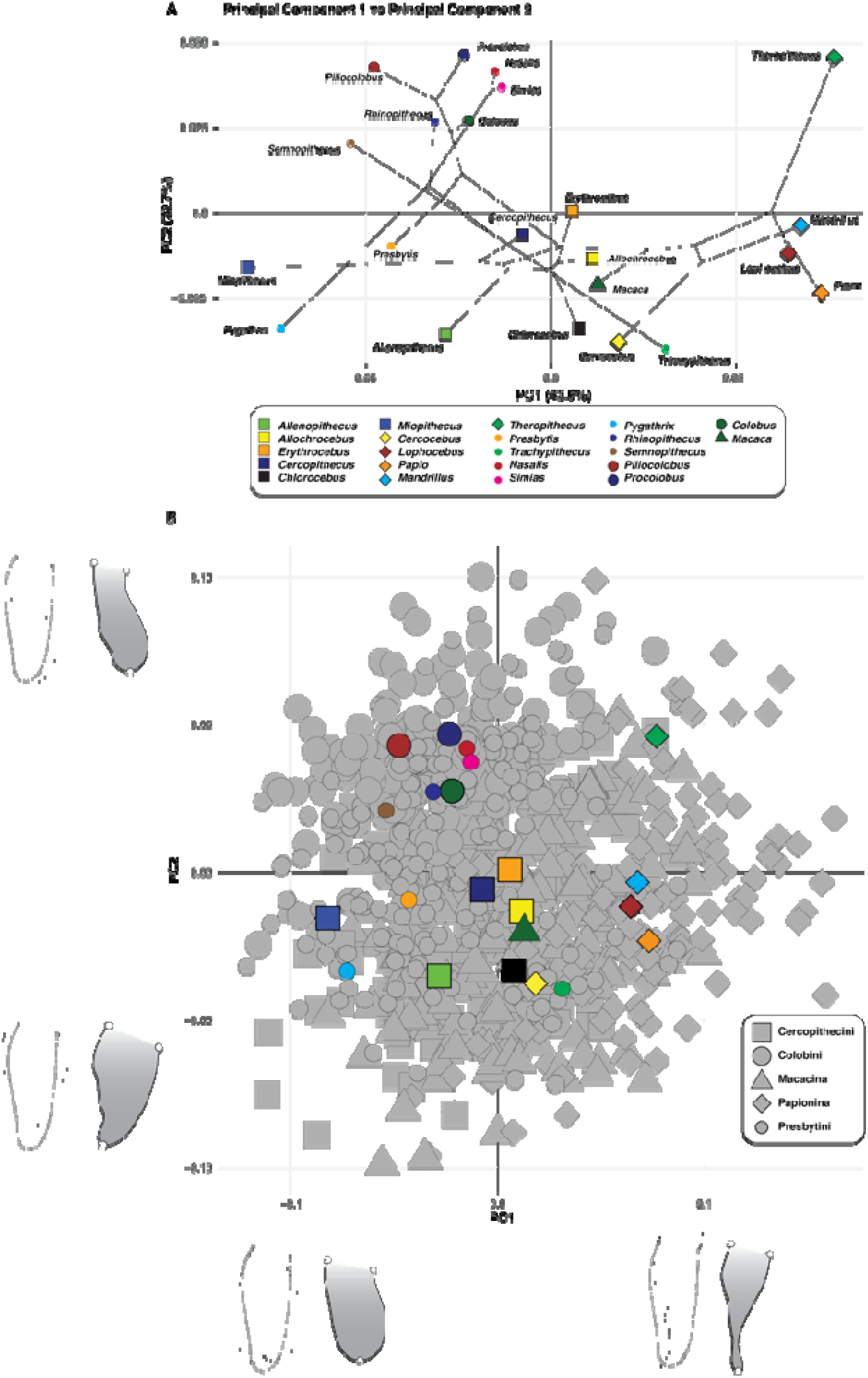
A) Phylomorphospace of PC1 and PC2 scores, and B) biplot of PC1 and PC2 scores. Corpus shape of specimens with maximum and minimum PC scores are shown along with shape changes associated with maximum and minimum PC scores.

Papionina, except for *Cercocebus*, presents highly positive scores on PC1*. Macaca* score is slightly lower than that of the Papionina on PC1 but is similar in value to that of *Cercocebus* (Figures 3 and 4A).

**Figure 4:**
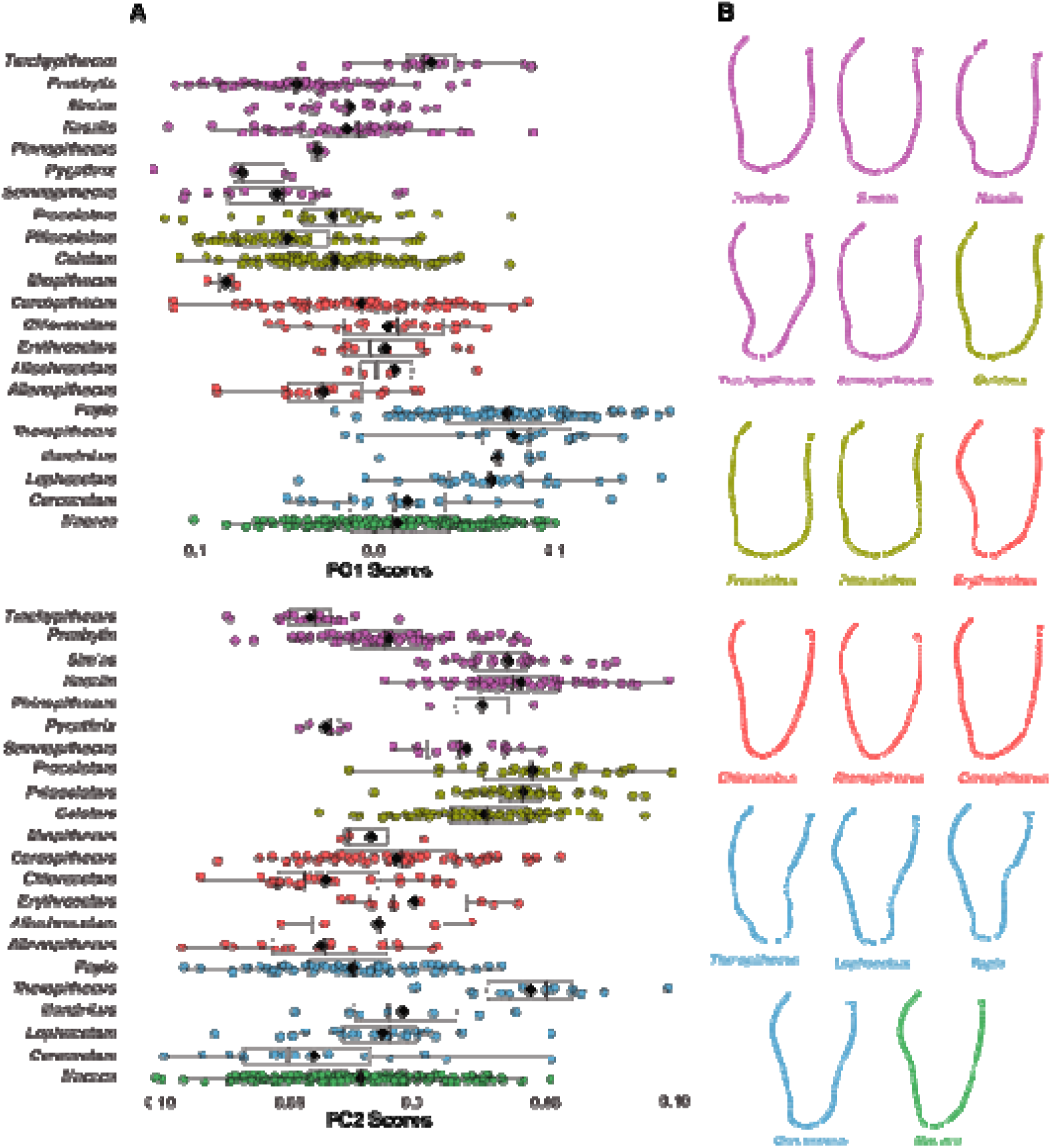
Boxplots of PC1 and PC2 scores with median (black line), mean (black diamond), third quartile and first quartile. Mean shape for taxa with n > 5 specimens and equally represented male and female specimens are figured on the right. For an extended version that includes taxa with n < 5 specimens and unequally represented male and female specimens (i.e., Pygathrix, Rhinopithecus, Allochrocebus, Miopithecus, and Mandrillus), see SOM Figure S4.

Among Cercopithecini, *Miopithecus* can be distinguished from all other guenon by showing a highly negative score on PC1. *Allenopithecus* also present a negative score compared to other guenons. *Cercopithecus*, *Chlorocebus*, and *Eythrocebus* present PC1 scores either slightly positive or slightly negative, hence clustering around zero. Among colobines, *Trachypithecus* stands apart by showing a positive PC1 score, aligning it with papionins. PC2, which accounts for ca. 23.0% of the variance, also permit to distinguish colobine from cercopithecines (Figure 3). Indeed, colobines exhibit positives scores while cercopithecines present negative ones. The only clear exceptions to this subfamilial pattern are *Pygathrix*, *Trachypithecus* and *Theropithecus*. Contrary to their closest relative, *Pygathrix* and *Trachypithecus* present negative scores while *Theropithecus* shows positive scores (Figure 4A). *Allenopithecus* and *Chlorocebus* also tend to be distinguished from their closest relatives on PC2, although differences in PC2 scores are less extreme than that of *Pygathrix*, *Trachypithecus*, and *Theropithecus*.

Shape changes along PC1 are driven by overall corpus robustness (Figure 3). Negative PC1 scores illustrates superoinferiorly short and buccolingually broad corpus, as exemplified by *Miopithecus* and *Pygathrix*. Positive PC1 scores depict superoinferiorly high and buccolingually thin (inferiorly) corpora. This morphology is typical of most papionins, which exhibit marked hollowing of the corpus at mid-height (i.e., corpus fossa). The convergence between *Trachypithecus* and papionins is explained by the extremely developed submandibular fossa of the former which engenders, as for corpus fossa, a thinning of the inferior aspect of the corpus (Figure 4B). PC2 shape changes are driven by the development of the lateral prominence, with positive PC2 scores exemplifying well-developed lateral prominences. High PC2 scores can be illustrated by the extremely developed lateral prominence of the African colobine *Procolobus* (Figure 3 and Figure 4B). In contrast, negative PC2 scores are not associated with lateral prominence but rather by a distal thinning of the corpus, as seen, for example, in *Trachypithecus*(Figure 4B).

### ALLOMETRY AND SIGNIFICANT DIFFERENCES IN PGLS AND PLS SLOPE VALUES

<u>Phylogenetic generalized least squares regressions</u>: Overall, PC1 scores are increasing with centroid size (Figure 5). Papionins present PC1 scores well above the cercopithecid regression line. This allometric relationship contrast with that of extant colobines, which present, apart from *Trachypithecus*, PC1 scores that fits below the cercopithecid regression line (Tables 1 and 2). Cercopithecini, except for *Allenopithecus* and *Miopithecus*, present PC1 scores above the cercopithecid regression line, similar to papionins (Figure 5). In summary, the corpus of papionins is high and thin in comparison with the broad and low corpus of colobines for a given range of size.

**Figure 5:**
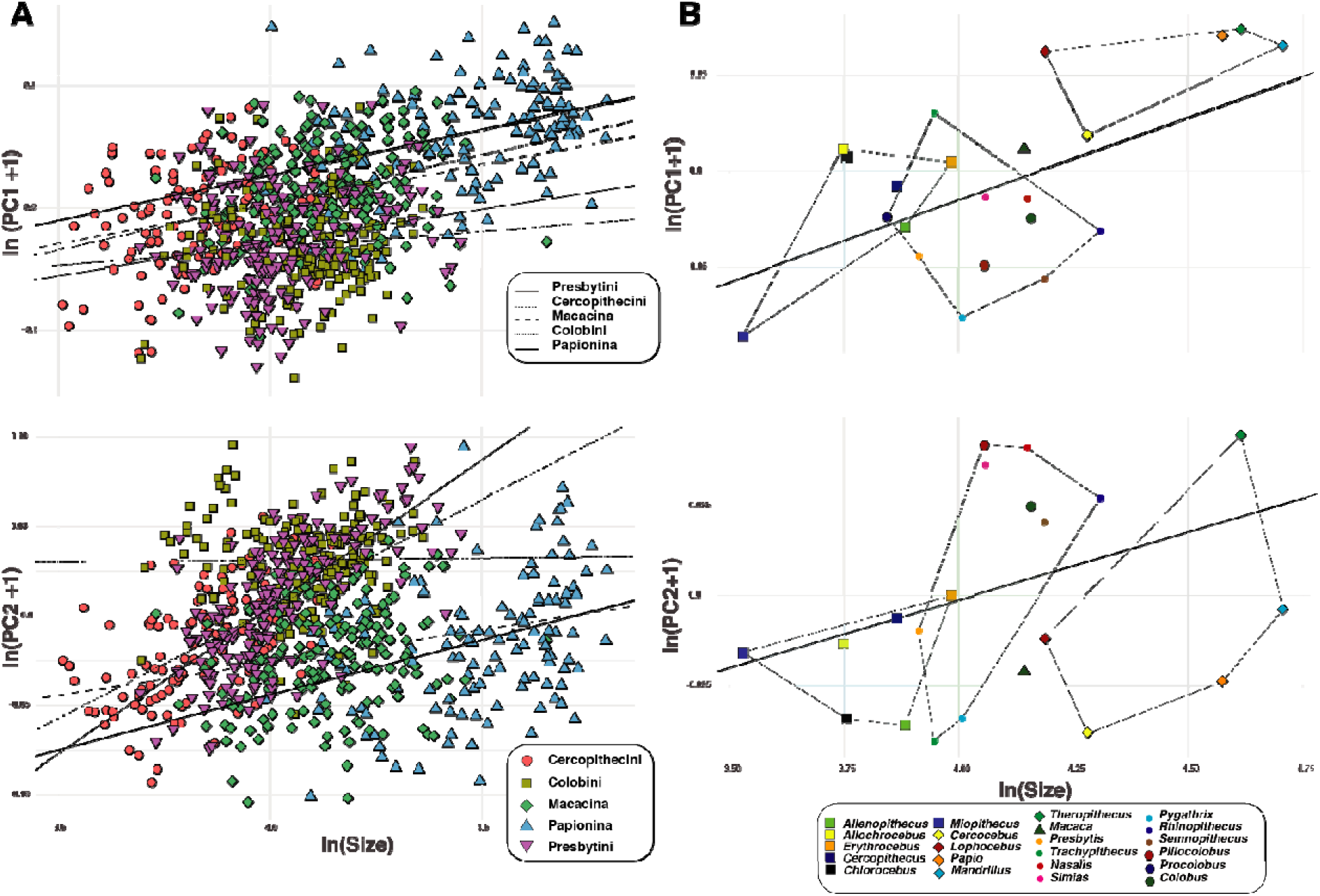
A) Biplot of the partial least squares regression of ln(PC scores +1) on ln(Centroid size), and B) Biplot of the phylogenetic generalized least squares regression of ln(PC scores +1) on ln(Centroid size).

**Table 1:** 95% confident interval of slope values of the PLS allometric regressions per subfamily, tribe, and subtribe.

| *Taxa* | **Slope values** |  |
| --- | --- | --- |
|  | *PC1* |  |
|  | 2.5 % | 97.5 % |
| *Cercopithecini* | 0.02 | 0.13 |
| *Colobini* | -0.01 | 0.07 |
| *Macacina* | 0.03 | 0.10 |
| *Papionina* | 0.04 | 0.10 |
| *Presbytini* | 0.01 | 0.09 |
| *Cercopithecinae* | 0.09 | 0.11 |
| *Colobinae* | < 0.01 | 0.07 |
|  | *PC2* |  |
|  | 2.5 % | 97.5 % |
| *Cercopithecini* | 0.09 | 0.15 |
| *Colobini* | -0.03 | 0.03 |
| *Macacina* | < 0.01 | 0.06 |
| *Papionina* | 0.03 | 0.09 |
| *Presbytini* | 0.14 | 0.19 |
| *Cercopithecinae* | 0.01 | 0.03 |
| *Colobinae* | 0.08 | 0.12 |

**Table 2:**
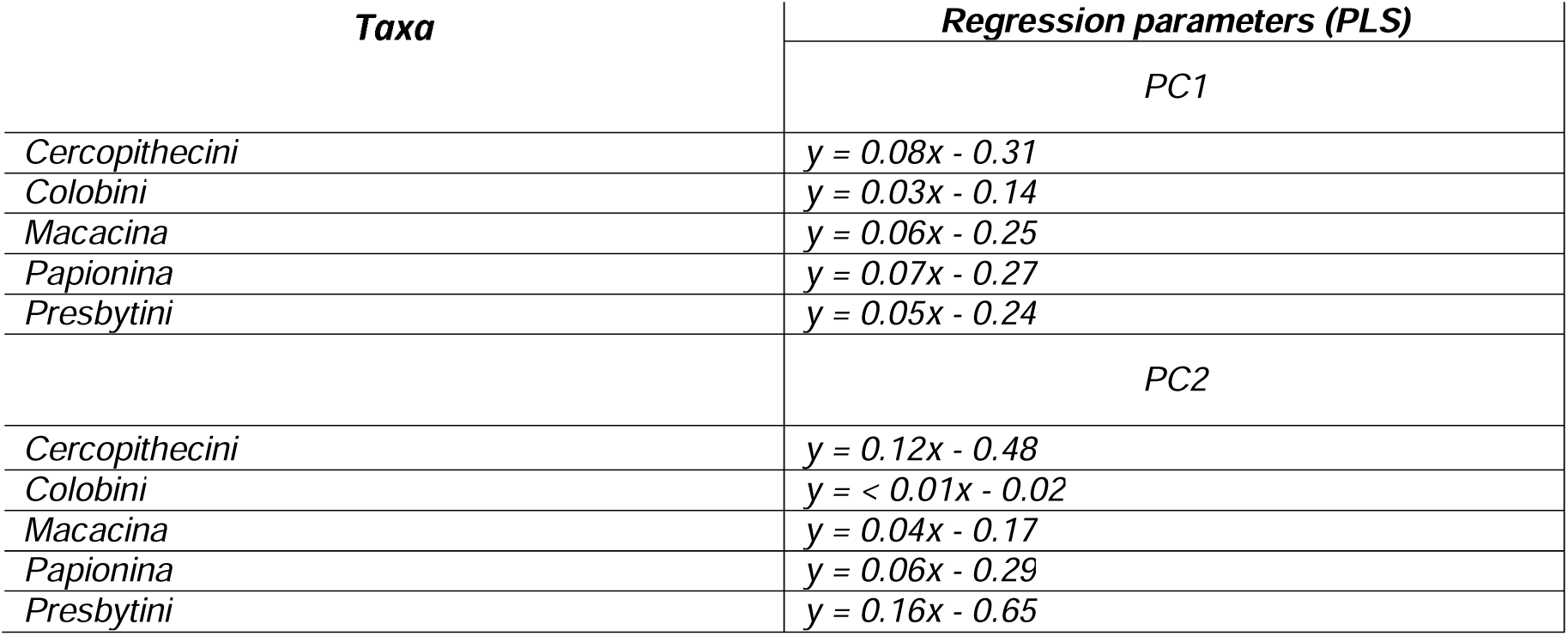
Parameters of the PLS allometric regressions per subfamily, tribe, and subtribe.

| *Taxa* | *Regression parameters (PLS)* |
| --- | --- |
|  | *PC1* |
| *Cercopithecini* | $y = 0.08x - 0.31$ |
| *Colobini* | $y = 0.03x - 0.14$ |
| *Macacina* | $y = 0.06x - 0.25$ |
| *Papionina* | $y = 0.07x - 0.27$ |
| *Presbytini* | $y = 0.05x - 0.24$ |
|  | *PC2* |
| *Cercopithecini* | $y = 0.12x - 0.48$ |
| *Colobini* | $y = < 0.01x - 0.02$ |
| *Macacina* | $y = 0.04x - 0.17$ |
| *Papionina* | $y = 0.06x - 0.29$ |
| *Presbytini* | $y = 0.16x - 0.65$ |

There is a positive allometry on PC2 scores, with scores increasing with centroid size (Figure 5; Tables 1 and 2). The allometric relationship of PC2 scores between subfamilies and tribes is identical to that describe for the PC2 axis of the PC1-PC2 biplot. In summary, *Theropithecus* present a high PC2 mean score compared to papionins with similar centroid size, the colobines *Trachypithecus*, *Pygathrix*, and *Presbytis* show low PC2 scores compared to similar-sized colobines, and *Cercopithecus*, and *Allenopithecus* have low PC2 scores compared to similar-sized guenons.

There is a highly significant difference in slope value between colobinae and cercopithecinae (Table 3) on the regression of PC1 and PC2 scores against centroid size. Apart from the pairs Cercopithecini - Macacina and Colobini - Presbytini, there is no significant differences in slope values between lower level taxa of each cercopithecid subfamily. Precisely, Presbytini shows a high slope value (a = 0.16, see Tables 1 and 3) compared to Colobini which present no relationship between PC2 scores and centroid size with a slope value of ca. 0 (Tables 1 and 3). Cercopithecini present a slope value much higher (a = 0.12, see Tables 1 and 3) than that of Macacina (a = 0.04, see Tables 1 and 3).

**Table 3:**
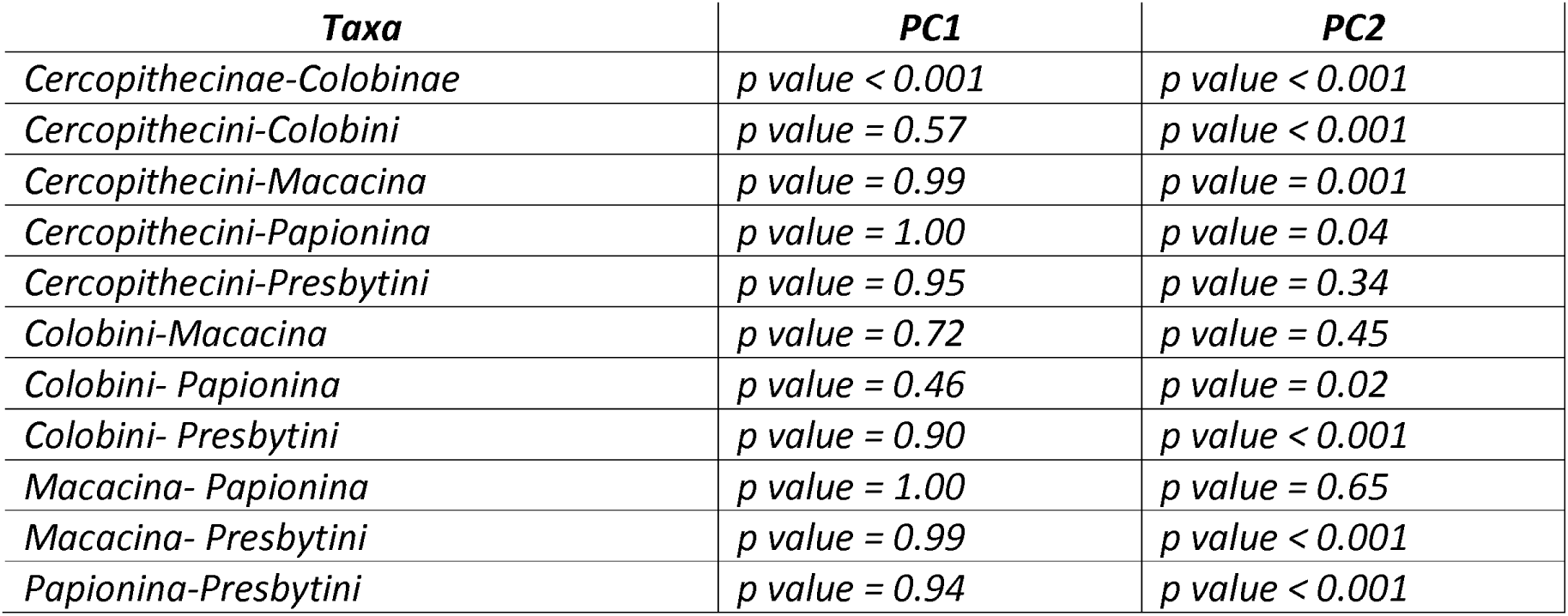
Differences in slope values (p-value) between subfamily, tribes and subtribes for the PLS regressions.

### SIGNIFICANT DIFFERENCES BETWEEN MALE AND FEMALE SPECIMENS (SEXUAL DIMORPHISM)

We detected significant differences in PC1 scores between male and female specimens of *Procolobus verus*, *Nasalis larvatus*, and *Papio anubis* (Table 4). In *P. anubis*, male specimens present higher scores relative to females whereas in *N. larvatus* and *Pro. verus*, females have lower scores relative to males (Figure 6A). The magnitude of differences is highly significant (*p* < 0.001) in *N. larvatus* and *P. anubis*, but significant (*p* < 0.05) in *Pro. verus* (Table 4). In sum, the corpus of female specimens from the sexually dimorphic taxa stated above is always more robust (i.e., having lower PC1 scores) than that of females (Figure 6B). This is well illustrated in comparing the mean shape of female and male *Procolobus verus* (Figure 6B).

**Figure 6:**
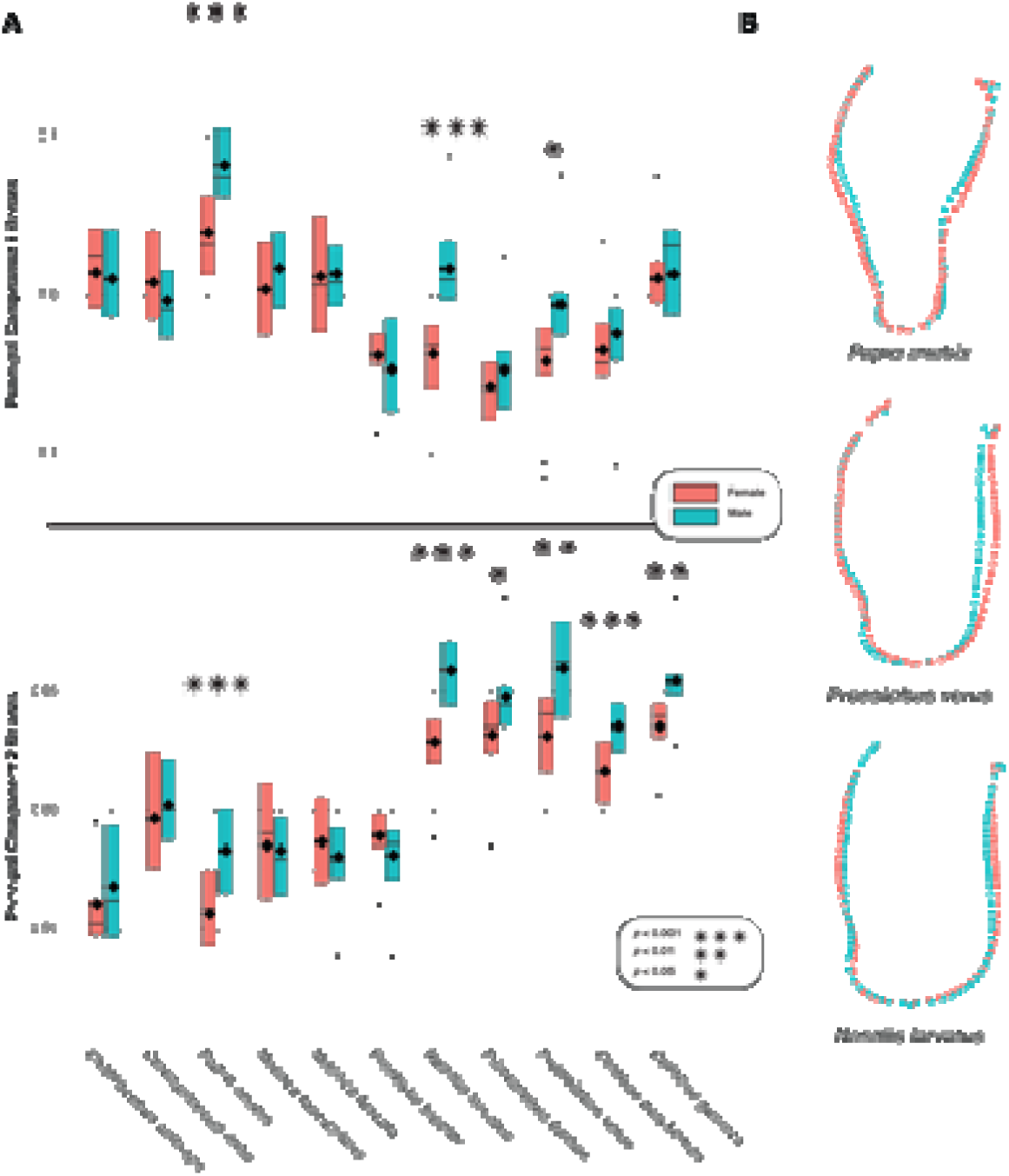
A) Boxplot of PC1 and PC2 scores of female and male specimens of selected cercopithecids, with median (black line), mean (black diamond), third quartile and first quartile. B) Mean corpus shape of female and male Papio anubis, Procolobus verus, and Nasalis larvatus.

**Table 4:** Significance and associated parameters of statistical models that test the effect of sexual dimorphism on principal component scores.

| **Formula** | **Test** | **p-value and test parameters** |
| --- | --- | --- |
| *Co. guereza* PC1 ~ Sex | ANOVA | $F = 0.04$<br>$p\text{-value} = 0.85$ |
| *Co. guereza* PC2 ~ Sex | | $F = 8.94$<br>$p\text{-value} = 0.007$ |
| *Co. polykomos* PC1 ~ Sex | | $F = 2.51$<br>$p\text{-value} = 0.12$ |
| *Co. polykomos* PC2 ~ Sex | | $F = 22.70$<br>$p\text{-value} < 0.001$ |
| *N. larvatus* PC1 ~ Sex | | $F = 41.70$<br>$p\text{-value} < 0.001$ |
| *N. larvatus* PC2 ~ Sex | | $F = 29.40$<br>$p\text{-value} < 0.001$ |
| *Pi. badius* PC1 ~ Sex | | $F = 0.80$<br>$p\text{-value} = 0.38$ |
| *Pi. badius* PC2 ~ Sex | | $F = 5.80$<br>$p\text{-value} = 0.02$ |
| *Pro. verus* PC1 ~ Sex | | $F = 5.93$<br>$p\text{-value} = 0.02$ |
| *Pro. verus* PC2 ~ Sex | | $F = 8.58$<br>$p\text{-value} = 0.007$ |
| *Pre. bicolor* PC1 ~ Sex | | $F = 0.33$<br>$p\text{-value} = 0.57$ |
| *Pre. bicolor* PC2 ~ Sex | | $F = 1.05$<br>$p\text{-value} = 0.32$ |
| *Cer. mitis* PC1 ~ Sex | | $F = 0.83$<br>$p\text{-value} = 0.37$ |
| *Cer. mitis* PC2 ~ Sex | | $F = 0.40$<br>$p\text{-value} = 0.53$ |
| *Ch. aethiops* PC1 ~ Sex | | $F = 0.07$<br>$p\text{-value} = 0.80$ |
| *Ch. aethiops* PC2 ~ Sex | | $F = 0.43$<br>$p\text{-value} = 0.52$ |
| *M. fascicularis* PC1 ~ Sex | | $F = 2.74$<br>$p\text{-value} = 0.10$ |
| *M. fascicularis* PC2 ~ Sex | ANOVA | $F = 0.09$<br>$p\text{-value} = 0.77$ |
| *M. fuscata* PC1 ~ Sex | | $F = 0.01$<br>$p\text{-value} = 0.91$ |
| *M. fuscata* PC2 ~ Sex | | $F = 0.49$<br>$p\text{-value} = 0.49$ |
| *P. anubis* PC1 ~ Sex | | $F = 16.80$<br>$p\text{-value} < 0.001$ |
| *P. anubis* PC2 ~ Sex | | $F = 15.00$<br>$p\text{-value} < 0.001$ |

A stronger sexually dimorphic signal is observed on PC2. Indeed, while three species were dimorphic on PC1 scores, six are dimorphic on PC2 scores. Precisely, highly significant differences (*p* < 0.001) are observed between male and female specimens of *Co. polykomos*, *N. larvatus*, and *P. anubis* (Table 4). Significant differences (*p* < 0.01) are observed between male and female specimens of *Co. guereza* and *Pro. verus*, but slightly less significant differences (*p* < 0.05) among *Pi. badius* (Figure 6A). In each dimorphic taxa, male specimens have higher PC2 scores relative to female. Morphologically, the corpus of male *N. larvatus* shows more developed lateral prominences and more excavated submandibular fossa (Figure 6B). The same is true for *Pro. verus*, where the male mean shape present a more developed submandibular fossa but differences in lateral prominences development is not as marked as in *N. larvatus* . In male *P. anubis*, the buccal mandibular fossa is more excavated in male than in females.

### SIGNIFICANT DIFFERENCES BETWEEN SUBFAMILIES AND TRIBES IN MULTIVARIATE DATA

A highly significant difference is seen on PC1 scores between colobines and cercopithecines (*p* < 0.001), with the former presenting significantly lower PC1 scores (Table 5). At tribal level, it is possible to significantly distinguish most pairs of tribes apart from the pairs Macacina - Papionina and Colobini-Presbytini.

**Table 5:**
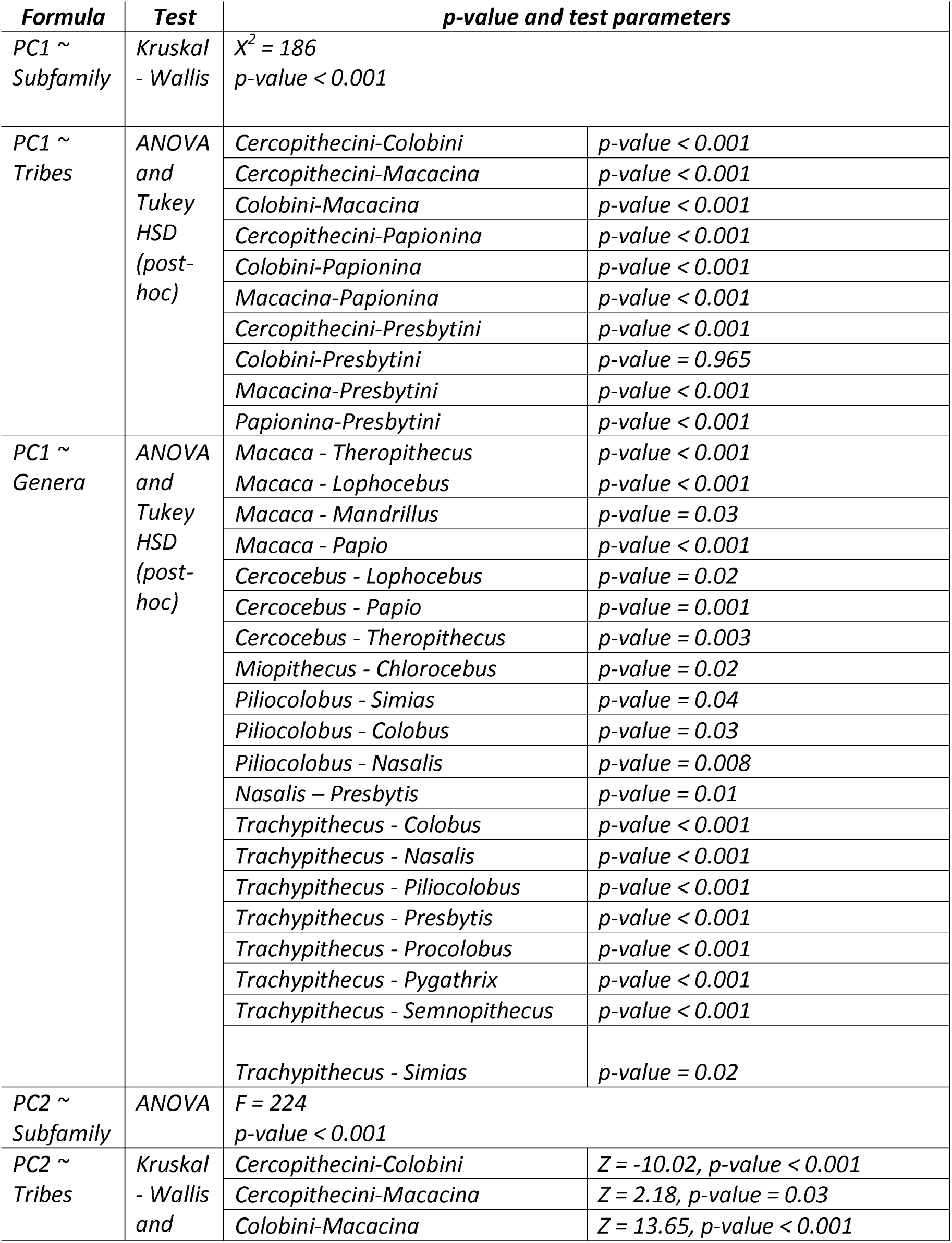
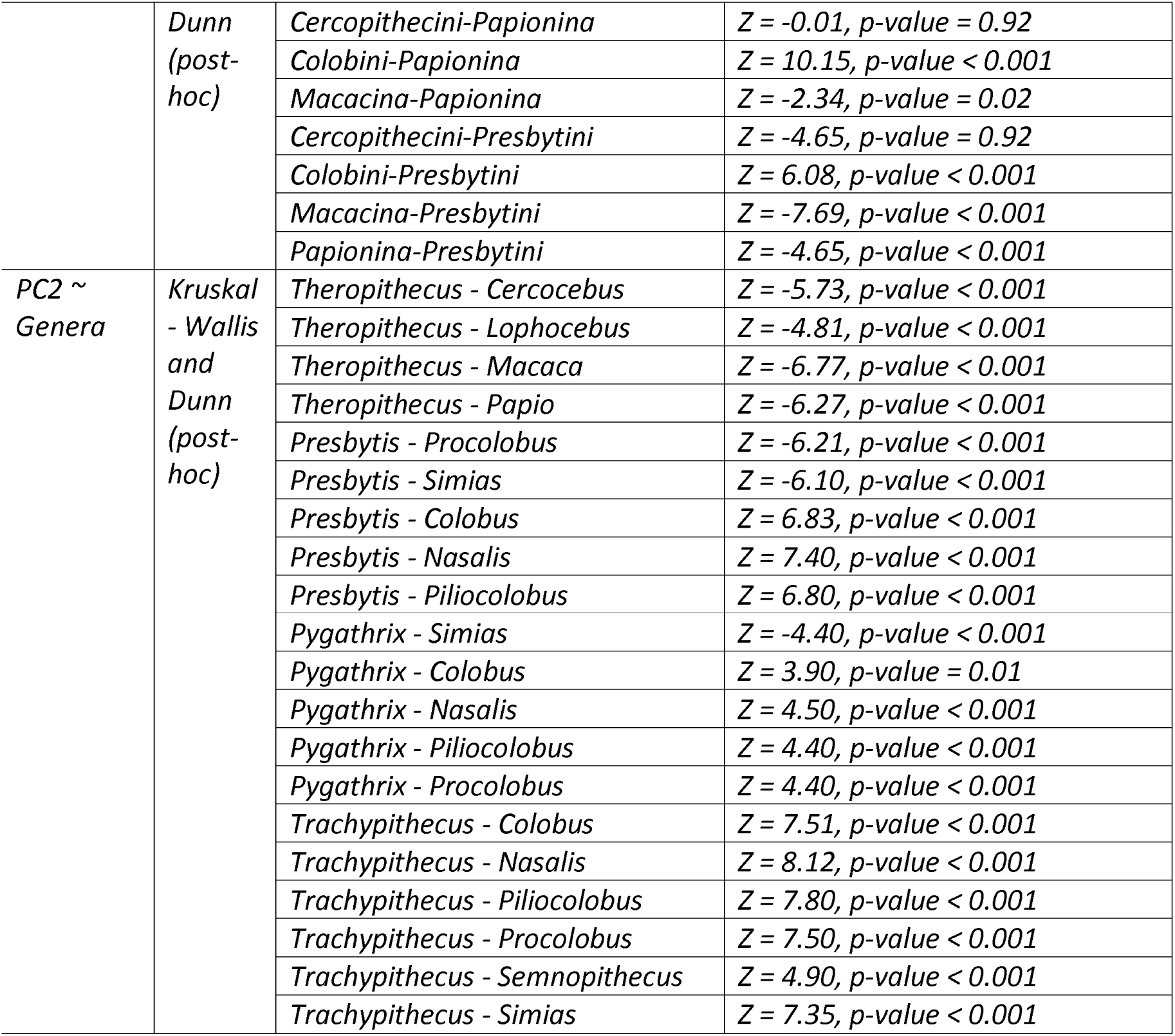
Significance and associated parameters of statistical models that test the effect of taxonomy on morphometric ratios.

Among colobines, *Piliocolobus*, *Nasalis* and *Trachypithecus* are highly distinctive on PC1. Indeed, *Trachypithecus* can be differentiated with a high level of confidence from most of the colobine taxa on the basis of its positive PC1 score (Table 5). *Nasalis* can also be significantly distinguished from *Presbytis* on PC1 by presenting a relatively higher score. Similarly, *Piliocolobus* is significantly different from *Colobus*, its sister taxa, on PC1. This differentiation is expressed by a lower PC1 score in *Piliocolobus* relative to *Colobus*. Cercopithecini are quite homogenuous in PC1 score and only the extremely negative score of *Miopithecus* can be distinguished from the positive score of *Chlorocebus* (Table 5). The *Macaca* corpus is distinct from that of other papionins. Indeed, its score on PC1 is significantly lower than that of all other papionins except *Cercocebus*. Among Papionina, *Cercocebus* is highly distinct from *Papio* and distinct from *Theropithecus* and *Lophocebus*.

Colobines can be singled out in a significant way from cercopithecines by presenting positive scores on PC2 (*p* < 0.001). All pairs of tribes can be significantly discriminated on PC2 apart from the pairs Cercopithecini-Presbytini and Cercopithecini-Papionina (Table 5).

At a genus level, the high PC2 scores of *Theropithecus* are significantly highly distinct from that of other Papionina and *Macaca* (Table 5). As for PC1, Cercopithecini are homogenuous on PC2 scores, with the pair *Cercopithecus*-*Chlorocebus* as the only one that is significantly different. Among colobines, *Presbytis* is significantly distinct from the Colobini, *Nasalis* and *Simias* on PC2 scores. The same pattern of difference is observed with *Pygathrix. Trachypithecus* is similarly distinct from the Colobini on PC2 scores, but it can also be significantly discriminated from *Nasalis*, *Simias*, and *Semnopithecus*.

### PHYLOGENETIC SIGNAL

PC1 scores follows a distribution expected under BM, as illustrated by its high Pagel’s λ value (Table 6). On the contrary, PC2 scores does not follow a distribution expected under BM and is hence not structured by phylogeny as demonstrated by its relatively low Pagel’s λ value (Table 6).

**Table 6:** Pagel’s λ parameters and significance of it.

|  | **Phylogenetic signal $\lambda$** |
| --- | --- |
| *PC1* | $\lambda = 1.00$, $p\text{-value} < 0.001$ |
| *PC2* | $\lambda = 0.21$, $p\text{-value} = 0.41$ |

## DISCUSSION

The aim of this study is to establish whether corpus anatomy is a reliable taxonomic discriminator and to explore sources of variation that significantly influence corpus shape. Using 2D geometric morphometric of corpus cross-sections set at M_1_-M_2_ junction, we demonstrated that cercopithecid subfamilies can be discriminated with a high level of statistical significance. Several significant differences between genera can be observed within each subfamily, in particular with the distinct corpus shape of *Trachypithecus* among colobines and that of *Theropithecus* and *Cercocebus* among Papionina. We also highlighted the presence of distinct allometric relationships between corpus shape and corpus centroid size between colobines and cercopithecines. Finally, we established that corpus shape is significantly influenced by sexual dimorphism in *P. anubis*, *N. larvatus* and *Pro. verus*. Overall, these results give a clearer picture of the anatomy of the corpus of extant cercopithecids and allow for discussion of several points regarding their taxonomy and ecology.

### Corpus shape, diet and dietary behaviors in cercopithecids

It has been hypothesized that deep and elliptical corpus were selected to counteract bending stresses set in the sagittal plane whereas broad and ovoid corpus were selected to resist transverse bending (wishboning) and axial torsion (Hylander, 1979; Smith, 1983; Daegling, 1989; Takahashi and Pan, 1994; Jablonski et al., 1998). These differences were contextualized in extant primates, and notably cercopithecids, with follivores having deep and elliptical corpora (Hylander, 1979; Bouvier, 1986; Ravosa, 1996; Jablonski et al., 1998, see also Takahashi and Pan, 1994) while primates feeding on mechanically challenging food such as hard-objects (e.g., seeds) or fibrous vegetation (e.g., grasses or seed pods) having broad and ovoid corpora (Hylander, 1979; Bouvier, 1986; Takahasi and Pan, 1994). The latter example raised the issue of equifinality of food that elicit high but singular loads, as opposed to food that requires low but repetitive loads (Daegling and Grine, 2017). This is particularly relevant when considering differences in dietary behaviors between colobines and cercopithecines, with the former chewing more often than the latter per ingestive event (McGraw et al., 2016; McGraw and Daegling, 2020). Misleading results have been found regarding ambiguous correlations between corpus shape and diet and/or dietary behaviors in extant cercopithecids (Smith, 1983; Daegling and McGraw, 2001; Daegling and Grine, 2017). As exposed in Smith (1983), discrepancies between corpus shape and dietary parameters result from the use of inaccurate diet and dietary behaviors categorization, inaccurate biomechanical proxies, and inaccurate biomechanical rationale for the interaction between corpus shape and mechanical forces (e.g., modelling the corpus as a hollow or enclosed beam and neglect of the role of the periodontal ligament in the distribution of bite forces). The latter is due in part to the lack of assessment of the internal bony structure of the corpus (Smith, 1983), which was subsequently greatly improved (Daegling, 1989; Daegling and Grine, 1991), but without providing clearer result as to the ecomorphological value of the internal structure of the corpus (Daegling and Grine, 1991; Daegling and Grine, 2007).

Although this paper does not seek to establish a link between any corpus shape and a particular function or diet, our findings are worth discussing in light of the ecomorphology of the cercopithecid mandible. It is a valid rationale to expect that diet and dietary behaviors are one of the main selective pressures shaping corpus anatomy, and the diverse diet and dietary behaviors of extant cercopithecids indeed provide an ideal framework for evaluating ecomorphological hypotheses (McGraw and Daegling, 2020). However, we demonstrated, on a large taxonomic scale, that the main axis of variation in our dataset (i.e., PC1 representing 43%) was primarily constrained by phylogeny, as previously suggested (Daegling, 2002; Daegling and Grine, 2007). Significant differences in PC1 scores were found between the major taxonomic groups: Colobinae, Cercopithecini, and Papionina. This result is not unexpected given the wide taxonomic range of our study and the relatively deep divergence times of the higher taxonomic ranks (e.g., ca. 17.6 Ma between Colobinae and Cercopithecinae) compared to the more recent radiation, for example, of papionins during the Pliocene. Phylogenetic inertia should therefore not be overlooked when inferring diet and / or dietary behaviors, as in some cases, it might blur the functional value of a trait. One interesting example drawn from our result is the close phenetic similarity between the sister taxa *Lophocebus* and *Papio* in spite of distinct diet and dietary behaviors. Indeed, both taxa present a distal thinning of the corpus, with fossae on the buccal aspect of the corpus and a large submandibular fossa on the lingual aspect. However, while *Lophocebus* is considered a seed eater, with a seed consumption frequency of up to 30% in *Lophocebus atterimus* (Horn, 1987) and *Lophocebus albigena* (Poulsen et al., 2001), seed consumption does not exceed 13% in *Papio anubis* (Okecha and Newton-Fisher, 2006; Johnson et al., 2012), not to mention that *Papio* has a more eclectic diet. Consequently, phylogenetic inertia, and thus the inherited corpus anatomy of the last common ancestor of *Papio* and *Lophocebus* may represent an adequate morphology to resist both the stresses induced by fruit incisal biting and molar crushing in *Lophocebus* (Happel, 1988) and the stresses induced by the more eclectic diet of *Papio* (Okecha and Newton-Fischer, 2006; Johnson et al., 2012). Given our methodological quantification of corpus shape, future ancestral character estimation and inclusion of fossil papionins, notably the fossil taxa *Papio robinsoni* (Freedman, 1957), *Papio hamadryas robinsoni* (Freedman, 1957) and *Lophocebus* cf. *albigena* (Jablonski et al., 2008b), could solve this issue and assert, with additional data from the fossil record, the hypothesis of phylogenetic inertia in the corpus shape of *Lophocebus* and *Papio* at M_1_/M_2_ junction.

### Corpus shape and allometry in cercopithecids

PGLS and GLS analyses yielded similar results, with cercopithecines of similar centroid size to colobines presenting much higher PC1 scores, reflecting their deeper corpus. Conversely, colobines, with the exception of *Trachypithecus*, have a more robust corpus. *Allenopithecus* and *Miopithecus* shows a more robust corpus compared to Cercopithecini of similar centroid size. Although this pattern may be influenced by a low sample size in *Miopithecus* (*n* = 3), the relatively large sample size of *Allenopithecus* (*n* = 16) allows us to conclude that the robust corpus of *Allenopithecus* does indeed deviate from the gracile corpus of other guenons.

In order to extract biomechanically relevant data, analyses interested in the scaling pattern of the catarrhine mandible regressed linear dimensions of the corpus and symphysis as a function of jaw length (Hylander, 1979; Bouvier, 1986; Ravosa, 1996; Vinyard and Ravosa, 1998; Jablonski et al., 1998; Daegling, 2001). Here, we regressed PC scores relative to the centroid size with the aim of using this protocol to include fossil specimens because, for the most part, the cercopithecid fossil record consists of fragmentary mandibles. Nevertheless, our results are consistent with previous analysis that have shown that the corpus of colobines is larger and more robust than that of cercopithecines (Bouvier, 1986; Ravosa, 1996).

### Corpus shape and sexual dimorphism in cercopithecids

Corpus depth and corpus gracility have been hypothesized to be a by-product of an elongated canine in sexually dimorphic primates (Chamberlain and Wood, 1985). The robustness of the corpus, represented by PC1 scores, is indeed highly dimorphic in *N. larvatus*, *Pro. verus* and *P. anubis* . These taxa are also documented as dimorphic in canine dimensions, and notably in crown height as demonstrated in Plavcan and Van Schaik (1992), and thus support a link between canine subocclusal anatomy and dimorphism. Similarly, the absence of sexual dimorphism on PC1 scores in *Colobus guereza* and *Presbytis bicolor* is consistent with the documented reduced sexual dimorphism of their canine dimensions (Lucas et al., 1986; Plavcan and Van Schaik, 1992; Hayes et al., 1995). However, the highly dimorphic lower canine of *Pi. badius* (Plavcan and van Schaick, 1992; Hayes et al., 1996), similar to its sister taxa *Pro. verus* (Hayes et al., 1996), does not appear to cause differences in corpus robustness on PC1. Indeed, no significant differences were detected in PC1 scores between male and female specimens of *Pi. badius*. Interestingly, while *Pro. verus* is highly dimorphic in canine dimensions, it is poorly dimorphic in molar dimensions (Hayes et al., 1996). Altogether, these results are difficult to reconcile and the *Pi. badius* vs. *Pro. verus* comparison is counterintuitive with regard to the effect of canine crown dimensions on corpus shape. In addition to this issue, we observed no difference in PC scores for male and female specimens of the dentally dimorphic species *Cer. mitis*, *Ch. aethiops*, *Ma. fuscata* and *Ma. fascicularis* (Plavcan and van Schaik, 1992), further blurring the link between subocclusal anatomy of the canine and dimorphism in corpus shape. Further work is needed to explain such results, including a quantification of the subocclusal anatomy of extant cercopithecids.

### The distinctive corpus shape of *Trachypithecus*, *Presbytis*, and *Pygathrix*

The corpus of *Trachypithecus* is significantly distinct from other colobines by presenting a distally thin corpus, most likely related to an enlargement of the submandibular fossa and reduced lateral prominences, as partly indicated by its score on PC2. The highly positive score of *Trachypithecus* on PC1 is also indicative of an elongated corpus. This result is in line with previous studies, which depicted the corpus of *Trachypithecus* as relatively deep relative to mandibular length (Bouvier, 1986; Wright et al., 2008). The corpus of *Trachypithecus* has been compared to that of *Pygathrix* as both are folivorous (Lippold, 1998; Zhou et al.,2006), but differ in their way of processing leaves (Wright et al., 2008). Although the study by Wright et al. (2008) suffers from a low sample size of *Trachypithecus* specimens (*n* = 4), our analysis, with a relatively much greater sample size (*n* = 18), yield a similar result (i.e., *Trachypithecus* having a deep corpus relative to *Pygathrix*), but with a finer resolution regarding corpus shape. Indeed, the corpus of *Pygathrix* is robust along its depth but present a deep, but restricted, submandibular fossa and a marked intertoral sulcus (SOM Figure S1) while that of *Trachypithecus* is thinning distally due to the presence of an enlarged submandibular fossa (Figure 4B). While *Pygathrix* possess a presaccus (gastric mill), and hence a quadripartite digestive system more efficient to process leaves and extract its nutrients (Caton, 1999; Wright et al., 2008; Matsuda et al., 2019), *Trachypithecus* lacks a presaccus (i.e., a tripartite stomach) and instead relies on intensive chewing and a larger molar area to fragment leaves and facilitate its digestion (Caton, 1999; Wright et al., 2008). Stomach morphology and physiology led Wright et al. (2008) to divide colobines into digestive folivores, comprising *Pygathrix*, *Nasalis*, *Rhinopithecus*, *Procolobus* and *Piliocolobus*, and ingestive folivores, comprising *Trachypithecus*, *Colobus*, *Semnopithecus*, and *Presbytis*. Although morphological differences between *Trachypithecus* and *Pygathrix* follow their distinct digestive physiology on PC1 scores, this is not true for the differences observed between two ingestive follivores (i.e., *Colobus* vs. *Trachypithecus* on PC1 and PC2 scores) or two digestive follivores (i.e., *Piliocolobus* vs. *Nasalis* on PC1 scores), demonstrating that corpus morphology is not a direct correlate of feeding behavior and digestive physiology.

The corpus shape of *Presbytis* is significantly distinct from that of other colobines, and in particular that of African colobines and *Nasalis*. It lacks enlarged lateral prominences and has a thin distal corpus contrary to the rounded corpus and large lateral prominences of African colobines and *Nasalis*. The precise phylogenetic relationships of *Presbytis* among the Asian colobines (Presbytini) are debated. *Presbytis* is alternatively interpreted as the sister taxon of *Trachypithecus* (Strasser and Delson, 1987; Sterner et al., 2006; Liedigk et al., 2012), as the sister taxon of the *Trachypithecus* / *Semnopithecus* group (Brandon-Jones, 1984), or as a basal Presbytini (Osterholz et al., 2008; Perelman et al., 2011). Our analysis shows that the corpus morphology of *Presbytis* is incompatible with that of *Trachypithecus* on the PC1 axis but not significantly distinct from it on PC2, sharing with the latter a distally thin corpus with weak lateral prominences. A close relationship between *Presbytis* and *Trachypithecus*, on the basis of corpus shape, is therefore ambiguous but is worth emphasizing the similarity between both taxa given their distally thin corpus. In addition, no significant differences were recorded between *Presbytis* and *Semnopithecus* on PC scores, a result consistent with a close relationship between *Presbytis*, *Semnopithecus* and *Trachypithecus*.

### A distinct corpus shape between *Piliocolobus* and *Colobus*

Daegling and McGraw (2001), based on the cross-sectional area of the corpus (computed using its maximum width and breadth) failed to distinguish *Co. polykomos* and *Pi. badius* . Our 2D morphometric geometric analysis gives another perspective to this previous result by demonstrating that, at a generic level, the corpus shape between *Colobus* and *Piliocolobus* is distinct. Compared to *Colobus*, *Piliocolobus* has a more robust corpus, as indicated by its relatively more negative PC1 scores. Although non-significant, the PC2 score of *Piliocolobus* also demonstrates its more developed lateral prominence compared to *Colobus*. In our study, the gracile corpus (and average positive score) of *Co. guereza* on PC1 leads the *Colobus* average score on the positive side, while *Co. polykomos* leads it on the negative side.

Due to its granivorous diet, Daegling and McGraw (2001) hypothesized that *Co. polykomos* had a more robust corpus in comparison to *Pi. badius* but failed to demonstrate it. We came to a similar conclusion as the interquartile range of PC1 scores overlaps greatly between *Pi. badius* and *Co. polykomos* . Future studies, focusing on interspecific differences, should provide additional data regarding the significance of this result.

The discovery of a distinct, quantifiable corpus shape between *Colobus* and *Piliocolobus* also paves the way for the identification of *Colobus* and *Piliocolobus* fossil remains. This is all the more relevant as the *Piliocolobus* fossil record is hitherto unknown. A future assessment of the Plio-Pleistocene fossil record could provide interesting results, particularly with regard to the initial attribution of *Colobus* specimens from Asbole (Frost and Alemseged, 2007) to *Piliocolobus* / *Procolobus* (Alemseged and Geraads, 2000) on the basis of a robust mandibular corpus.

### Similarities in corpus shape between *Colobus* and *Nasalis*

*Colobus* and *Nasalis* share similar scores on the corpus shape PC1-PC2 biplot shape despite having distinct symphyseal anatomy and dental dimensions. A more developed inferior torus, according to Pallas et al. (2019), is observed in *Nasalis*, and a broader M_1_ and a longer M_3_ according to the Presbytini dental pattern identified in Pan et al. (2004). Differences in digestive physiology (i.e., quadripartite stomach morphology in *Nasalis* vs. tripartite stomach morphology in *Colobus*; Matsuda et al., 2019), feeding behaviors and diet (i.e., granivory is more typical of *Co. polykomos*), in addition to a deep divergence time (ca. 12.30 Ma), make this morphological convergence unexpected. First, it illustrates the fact that, in *Nasalis* and *Colobus*, symphyseal and corpus shape are decoupled and influenced by distinct selective pressures. Second, it illustrates, once again, the difficulty of linking morphologies to a particular feeding behavior, diet, or physiology.

### The distinct corpus of *Macaca*, *Theropithecus*, and *Cercocebus*

Within Papionini, *Macaca* can be distinguished from all other genera, with the exception of *Cercocebus*, by presenting a low PC1 scores reflecting its more robust corpus. This result highlights the distinctiveness of the *Macaca* corpus, as previously suggested based on the absence of corpus fossa in extant and fossil *Macaca* representatives (Delson, 1980; Gilbert, 2007, Harrison, 2011). According to cladistic analyses, *Macaca* is generally considered to be a plesiomorphic papionin (Gilbert, 2013; Pugh and Gilbert, 2018), lacking some of the cranial and mandibular distinctive traits of extant Papionina (e.g., anteorbital drop).

Our *Macaca* sample is dominated by *M. fascicularis* (*n* = 99) and *M. fuscata* (*n* = 30), which belong to the same clade, the *M. fascicularis* group (Fooden, 1979; Delson, 1980; Abegg and Thierry, 2002), with limited specimens of the clades *M. silenus* / *M. sylvanus* (n = 10), *M. sinica* (n = 14; including *M. radiata*, *M. assamensis*, and *M. thibetana*), and *M. arctoides* (n = 5). The phylogenetic position of *M. sylvanus*, *M. arctoides*, and *M. nigra*, remains controversial, especially in regard to their affinity with the *M. sinica* and *M. fascicularis* clades or their placement into their own phyletic lineages (Fooden, 1976; Delson et al. 1980; Abegg and Thierry, 2002; Li et al., 2009). The inclusion of additional specimens from the latter taxa has the potential to refine our results regarding the variability of the *Macaca* corpus and to raise new hypotheses regarding phylogenetic relationships inside the *Macaca* clade.

*Theropithecus gelada* is the sister taxon of the *Lophocebus* / *Papio* clade, and is the only remaining species of the successful adaptive radiation of *Theropithecus* in the Plio-Pleistocene (Leakey, 1993). Extant geladas are mostly restricted to the mountainous areas of the Central Ethiopian Plateau and have a specific diet consisting mainly of grasses (Bergman and Beehner, 2013; Fashing and Nguyen, 2016). On the main axis of variation (i.e., PC1 scores), *Th. gelada* conforms to the shape of *Papio* and *Lophocebus* (i.e., deep corpus with the presence of mandibular fossae) but it is significantly distinct from them on PC2. This distinction illustrates the distally everted (curved buccally in the coronal plane) corpus of *Th. gelada* which reflects the developed lateral prominence of some colobines on PC2. The distal part of the corpus is also much more robust in *Th. gelada* than in *Papio*, reflecting the greater extension of the inferior transverse torus (ITT) in *Th. gelada* (Dechow and Singer, 1984; Delson and Dean, 1993). This observation is also consistent with the emended diagnosis of the genus *Theropithecus* which characterized the corpus of *Theropithecus* as presenting ’buttressed anterior inferior margins’ according to Getahun et al. (2023:7). The functional implication of an everted corpus in *Th. gelada* is unknown but a link with the development of the ITT is consistent with our data. Qualitative differences in the development (depth) of the corpus fossae development was hypothesized to distinguish subspecies of the *Theropithecus oswaldi*lineage (*T. o. darti*, *T. o. oswaldi* and *T. o. leakeyi*) from *T. brumpti* and *T. gelada* (Eck, 1993; Leakey, 1993; Frost and Delson, 2002; Frost, 2007; Frost and Alemseged, 2007; Frost et al., 2014, 2020; Getahun et al., 2023). The morphometric study of Dechow and Singer (1984), using linear corpus dimensions, demonstrated that this trait is not taxonomically reliable to distinguish *Theropithecus* fossil specimens from Makapan and Hopefield (South Africa), is influenced by sexual dimorphism and shows great intraspecific and interspecific variability in extant papionins. Our data concur with these results by demonstrating no significant difference in PC1 scores between the large papionins *Papio*, *Theropithecus* and *Mandrillus* (but see the paragraph below), in addition that corpus fossae are indeed influenced by sexual dimorphism in *Papio anubis*. Our quantification methods will benefit to the study of the *Theropithecus* fossil record, especially for *T. o. darti* and *T. o. oswaldi*, where corpus fossae variable in depth were described on qualitative basis only (Frost and Delson, 2002; Frost et al., 2020; Getahun et al., 2023).

*Cercocebus* does not cluster with African papionins on PC1 scores, the main axis of variation, and is significantly distinct from *Papio*, *Lophocebus* and *Theropithecus*. Such differences express a robust and shorter (superoinferiorly) corpus in *Cercocebus* relative to *Papio*, *Lophocebus* and *Theropithecus*. A deep corpus at the M_1_ was considered a synapomorphic feature of the African papionins according to Gilbert’s (2013) cladistic analysis. Under this phylogenetic hypothesis, the close similarity between *Cercocebus* and *Macaca* is intriguing, and may illustrate a morphological convergence due to shared selective pressures or alternatively, the retention of the plesiomorphic corpus shape condition in both taxa. Differences in analytical protocol should also be considered as our outline analysis differs in methodology from linear dimensions (Gilbert, 2013). Shallow corpus fossae, especially in female specimens, has also been highlighted as a synapomorphy of the *Cercocebus* / *Mandrillus* clade (Gilbert, 2007; Harrison, 2011). Our data confirm this hypothesis for *Cercocebus* but not for *Mandrillus*, but this discrepancy can be explained by the absence of female *Mandrillus* in our sample.

## CONCLUSION

We have demonstrated that the transverse cross-section of corpus shape at the M_1_-M_2_ junction effectively discriminates extant cercopithecid taxa at different taxonomic levels. We found that the main axis of variation of the corpus shape is structured following the distribution expected under Brownian motion, demonstrating its strong phylogenetic constraint. On a subfamilial level, we highlighted the distinctiveness of the robust colobine corpus in regard to the more gracile corpus of cercopithecines. On a generic level, the corpus shape of *Trachypithecus* converges with that of the papionins and is extremely different from that of other colobines, with slight lateral prominences and large submandibular fossae that result in an inferior tapering of the corpus. *Presbytis* and *Pygathrix* also present slight lateral prominences compared to other colobines. The corpus of *Allenopithecus* is more robust compared to other guenons and converges with that of the colobines on the main axis of variation (i.e., PC1). *Macaca* has a corpus distinct from that of the African papionins by the absence of mandibular fossa. Surprisingly, *Cercocebus* shows a corpus shape convergent with that of *Macaca* and is significantly distinguished from African papionins by shallow mandibular fossae. *Theropithecus* exhibits a corpus shape consistent with that of its closest relatives on PC1, but differs on PC2, and features a distally everted corpus that contrasts with the straight corpus of *Papio* and *Lophocebus*. In addition to taxonomy, sexual dimorphism also influences corpus shape, particularly in *Papio anubis*, *Nasalis larvatus*, and *Procolobus verus* . Variation in overall robustness, and in the development of the lateral prominences and submandibular fossae explains the variation due to sexual dimorphism in *Pro. verus* and *N. larvatus*, with females having more robust corpus but males showing more developed submandibular fossae and, in *N. larvatus*, more pronounced lateral prominences. Mandibular fossae are also dimorphic in *P. anubis*, with males having deeper fossae. Our data do not clearly support the hypothesis of an influence of dental subocclusal anatomy, and in particular canine subocclusal anatomy, in shaping corpus differences between males and females. This is exemplified by the absence of sexual dimorphism on PC1 scores in *Pi. badius* but the presence of dimorphism in *Pro. verus*, although both taxa have sexually dimorphic lower canine crown dimensions. Further work is needed to understand the proximate driver of intraspecific corpus shape variation in colobines. Our study proposes a new approach to studying the corpus anatomy of extant and fossil cercopithecids. It should benefit greatly in the future from the inclusion of fossil taxa and a more thorough assessment of taxonomic variation at the species level.

## ACKNOWLEDGEMENTS

The authors especially acknowledge the support of curators and staff of the National Museums of Kenya (Osteology Division, Nairobi), the Royal Museum for Central Africa (Dr. Emmanuel Gilissen and Mathys Rotonda from the Division of Mammals, Tervuren), the Center for the Evolutionary Origins of Human Behavior (Assoc. Pro. Takeshi Nishimura the Division of Mammals), the US National Museum of Natural History (D. Schlitter, Division of Mammals), and the Bavarian State Collection for Zoology (Dr. Anneke van Heteren and Michael Hiermeier) for the study of skeletal collections under their care. We also thank the NSF BCS 1552848 attributed to Dr. D. Boyer, and Assoc. Prof. Claire Terhune (University of Arkansas) for data sharing via the MorphoSource platform. We thank the Domestic Research Program of Ryokoku University. This research was financially supported by the Japan Society for the Promotion of Science JSPS Kakenhi (23H02562 and 16H02757) to MN.

## AUTHOR CONTRIBUTIONS

L.P. M.N. and Y.K. designed the study, L.P. acquired the data, L.P. M.N. and Y.K. interpreted data, L.P. drafted the manuscript, L.P., M.N. and Y.K. critically revised and approved the manuscript.

