## Supplementary material for "Anatomy of the mandibular corpus of extant cercopithecids : taxonomy and variation": Su: Pallas et al.,2023_Corpus_Supplementary.docx

SOM Figure S1: Outline of the corpus of *Pygathrix nemaeus* at M_1_-M_2_ junction

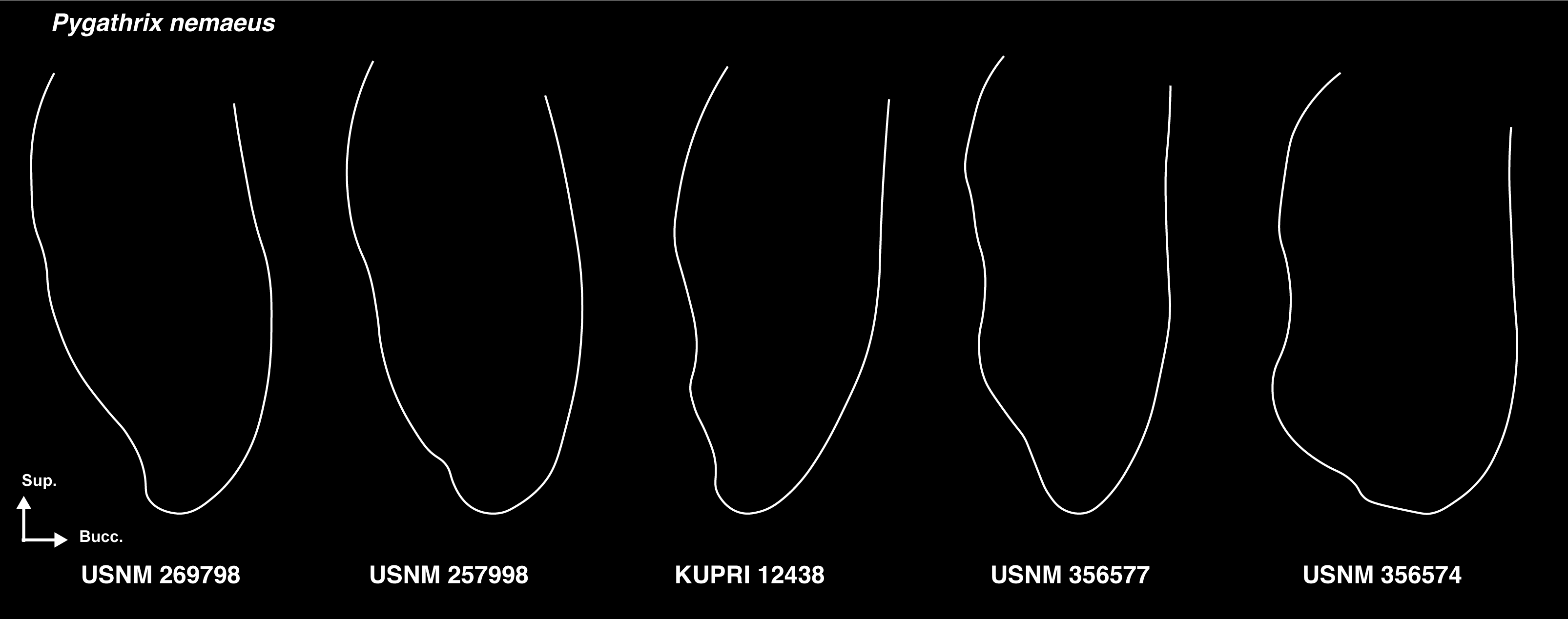

SOM Table S2: Normality and homoscedasticity of the model residuals that test for the effect of taxonomy on principal component scores.

| Formula | Normality (Shapiro-Wilk) | Homoscedasticity (Bartlett) |
| --- | --- | --- |
| PC1 ~ Subfamily | W = 1  *p*-value = 0.05 | K^2^ = 25  *p*-value < 0.001 |
| PC1 ~ Tribes | W = 1  *p*-value = 0.7 | K^2^ = 9  *p*-value = 0.06 |
| PC1 ~ Genera | W = 1  *p*-value = 0.6 | K^2^ = 27  *p*-value = 0.2 |
| PC2 ~ Subfamily | W = 1  *p*-value = 0.1 | K^2^ = 0.2  *p*-value = 0.6 |
| PC2 ~ Tribes | W = 1  *p*-value = 0.4 | K^2^ = 36  *p*-value < 0.001 |
| PC2 ~ Genera | W = 1  *p*-value = 0.7 | K^2^ = 46  *p*-value = 0.001 |

SOM Table S3: Normality and homoscedasticity of the model residuals that test for the effect of sex on morphometric ratios.

| Formula | Normality (Shapiro-Wilk) | Homoscedasticity (Bartlett) |
| --- | --- | --- |
| *Co. guereza* PC1 ~ Sex | W = 1  *p*-value = 0.9 | K^2^ = 0.5  *p*-value = 0.5 |
| *Co. polykomos* PC1 ~ Sex | W = 1  *p*-value = 0.6 | K^2^ = 0.6  *p*-value = 0.4 |
| *N. larvatus* PC1 ~ Sex | W = 1  *p*-value = 0.7 | K^2^ = 0.3  *p*-value = 0.6 |
| *Pi. badius* PC1 ~ Sex | W = 1  *p*-value = 0.4 | K^2^ = 0.2  *p*-value = 0.6 |
| *Pro. verus* PC1 ~ Sex | W = 1  *p*-value = 0.6 | K^2^ = 0.05  *p*-value = 0.8 |
| *Pre. bicolor* PC1 ~ Sex | W = 1  *p*-value = 0.5 | K^2^ = 4  *p*-value = 0.04 |
| *Ce. mitis* PC1 ~ Sex | W = 1  *p*-value = 1 | K^2^ = 0.7  *p*-value = 0.4 |
| *Ch. aethiops* PC1 ~ Sex | W = 1  *p*-value = 0.4 | K^2^ = 0.1  *p*-value = 0.7 |
| *M. fascicularis* PC1 ~ Sex | W = 1  *p*-value = 0.8 | K^2^ = 1  *p*-value = 0.3 |
| *M. fuscata* PC1 ~ Sex | W = 1  *p*-value = 0.2 | K^2^ = 2  *p*-value = 0.2 |
| *P. anubis* PC1 ~ Sex | W = 1  *p*-value = 0.2 | K^2^ = 0.05  *p*-value = 0.8 |
| *Co. guereza* PC2 ~ Sex | W = 0.9  *p*-value = 0.2 | K^2^ = 0.3  *p*-value = 0.6 |
| *Co. polykomos* PC2 ~ Sex | W = 1  *p*-value = 0.6 | K^2^ = 0.2  *p*-value = 0.7 |
| *N. larvatus* PC2 ~ Sex | W = 1  *p*-value = 0.8 | K^2^ = 0.6  *p*-value = 0.4 |
| *Pi. badius* PC2 ~ Sex | W = 1  *p*-value = 0.8 | K^2^ = 0.04  *p*-value = 0.8 |
| *Pro. verus* PC2 ~ Sex | W = 1  *p*-value = 0.5 | K^2^ = 0.005  *p*-value = 0.9 |
| *Pre. bicolor* PC2 ~ Sex | W = 0.9  *p*-value = 0.2 | K^2^ = 1  *p*-value = 0.2 |
| *Ce. mitis* PC2 ~ Sex | W = 1  *p*-value = 0.3 | K^2^ = 0.06  *p*-value = 0.8 |
| *Ch. aethiops* PC2 ~ Sex | W = 0.9  *p*-value = 0.07 | K^2^ = 2  *p*-value = 0.2 |
| *M. fascicularis* PC2 ~ Sex | W = 1  *p*-value = 0.7 | K^2^ = 3  *p*-value = 0.09 |
| *M. fuscata* PC2 ~ Sex | W = 1  *p*-value = 0.8 | K^2^ = 3  *p*-value = 0.1 |
| *P. anubis* PC2 ~ Sex | W = 1  *p*-value = 0.6 | K^2^ = 0.004  *p*-value = 1 |

SOM Figure S4: Mean shape of the corpus of extant cercopithecids, including *Pygathrix*, *Rhinopithecus*, *Allochrocebus*, *Miopithecus*, and *Mandrillus*.

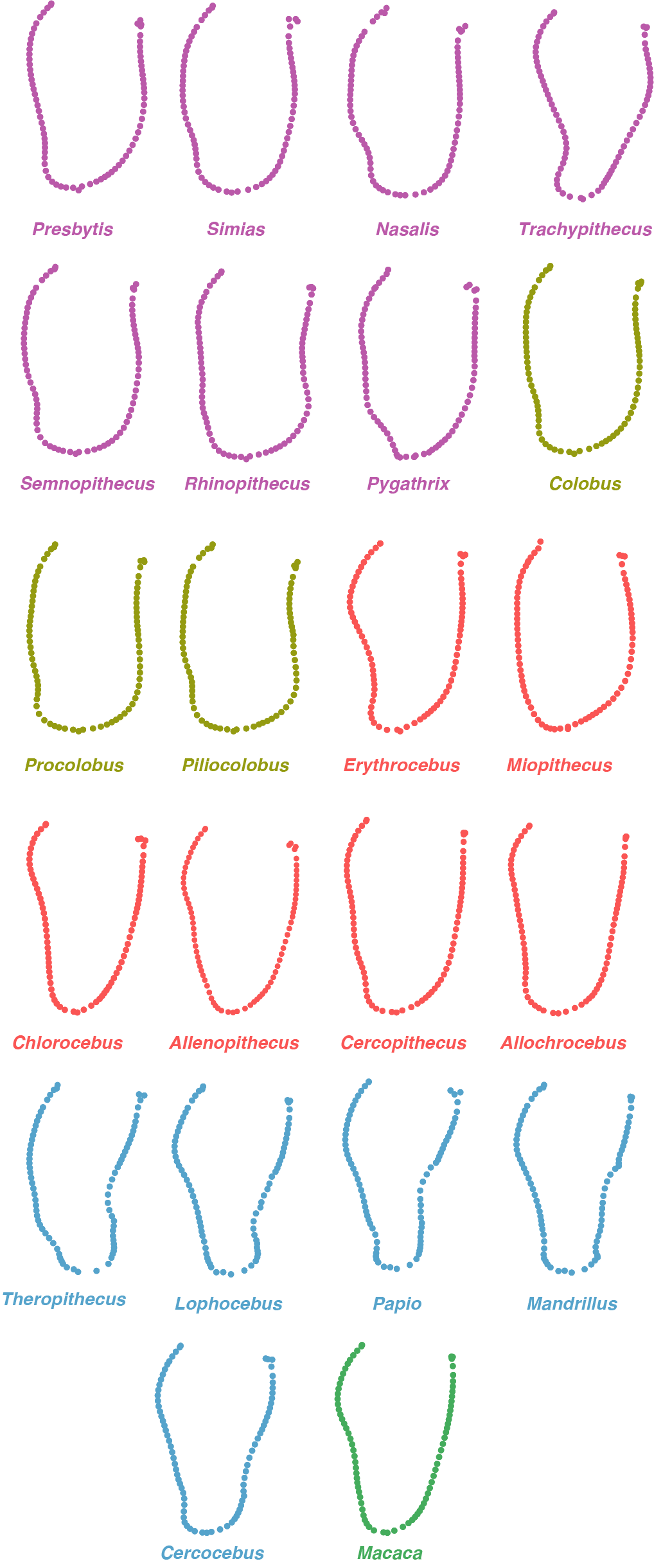
