## Supplementary figures and images for "Anatomy of the mandibular corpus of extant cercopithecids : taxonomy and variation"

### Figure_1.png

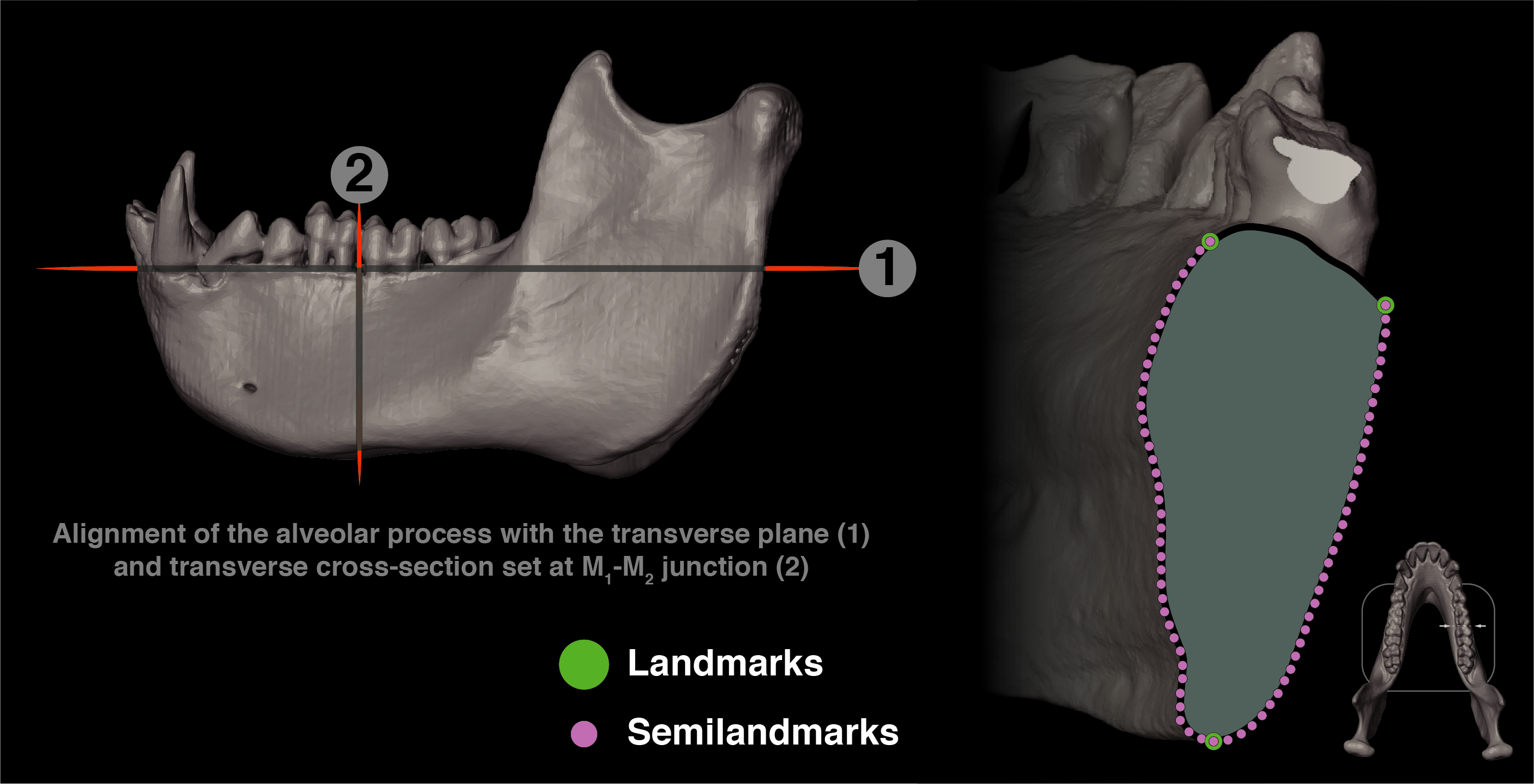

### Figure_2.png

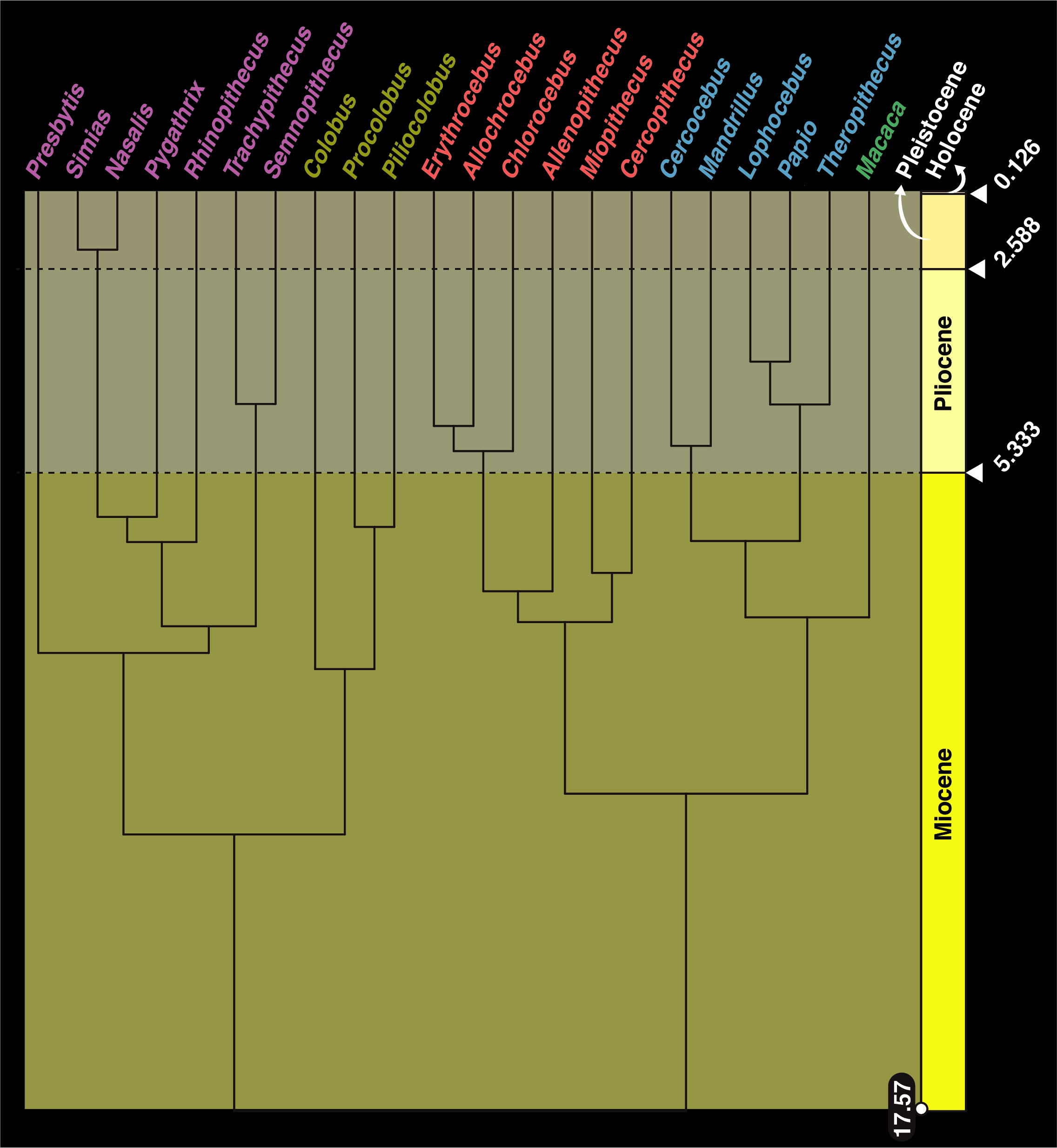

### Figure_3.png

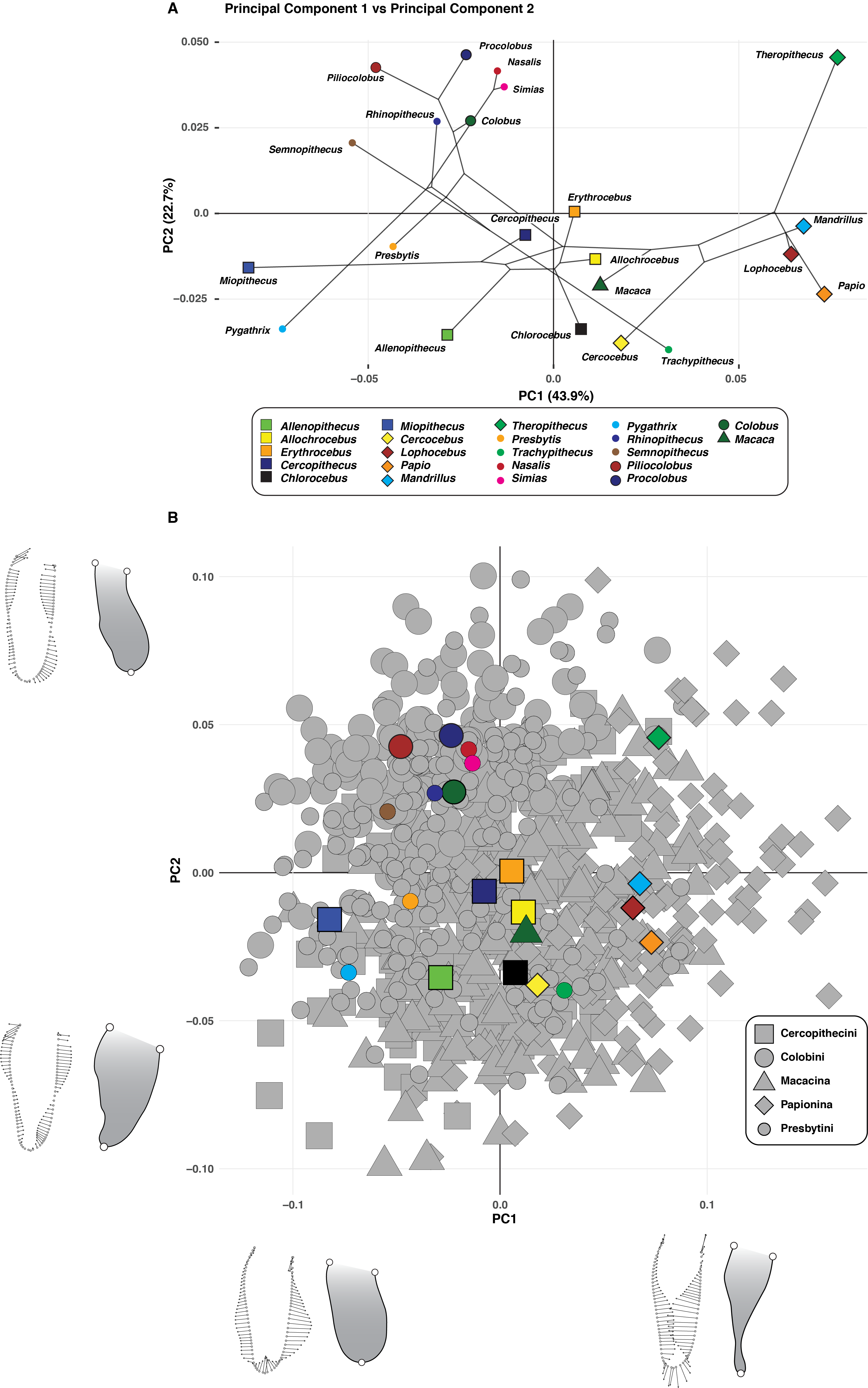

### Figure_4.png

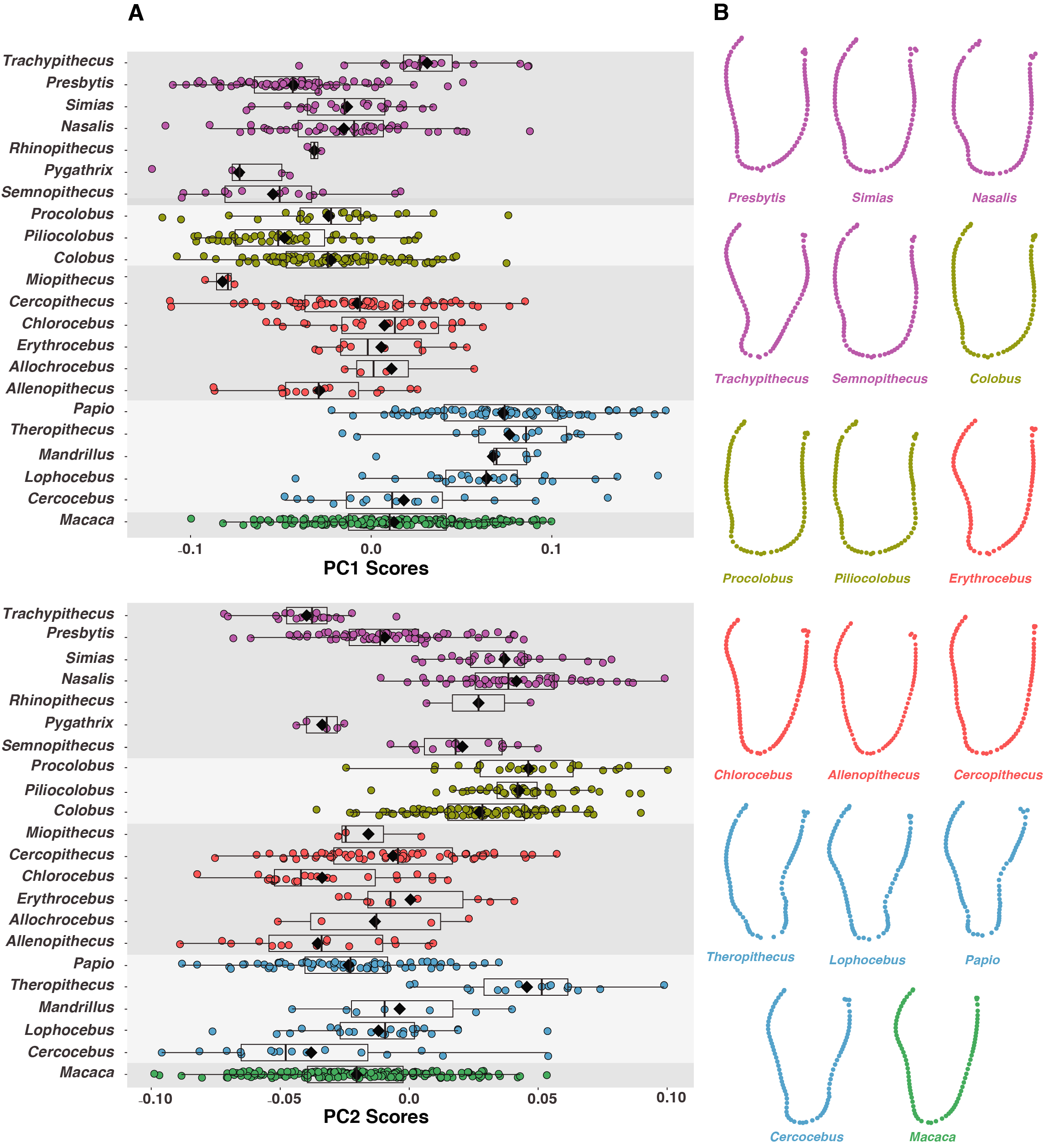

### Figure_5.png

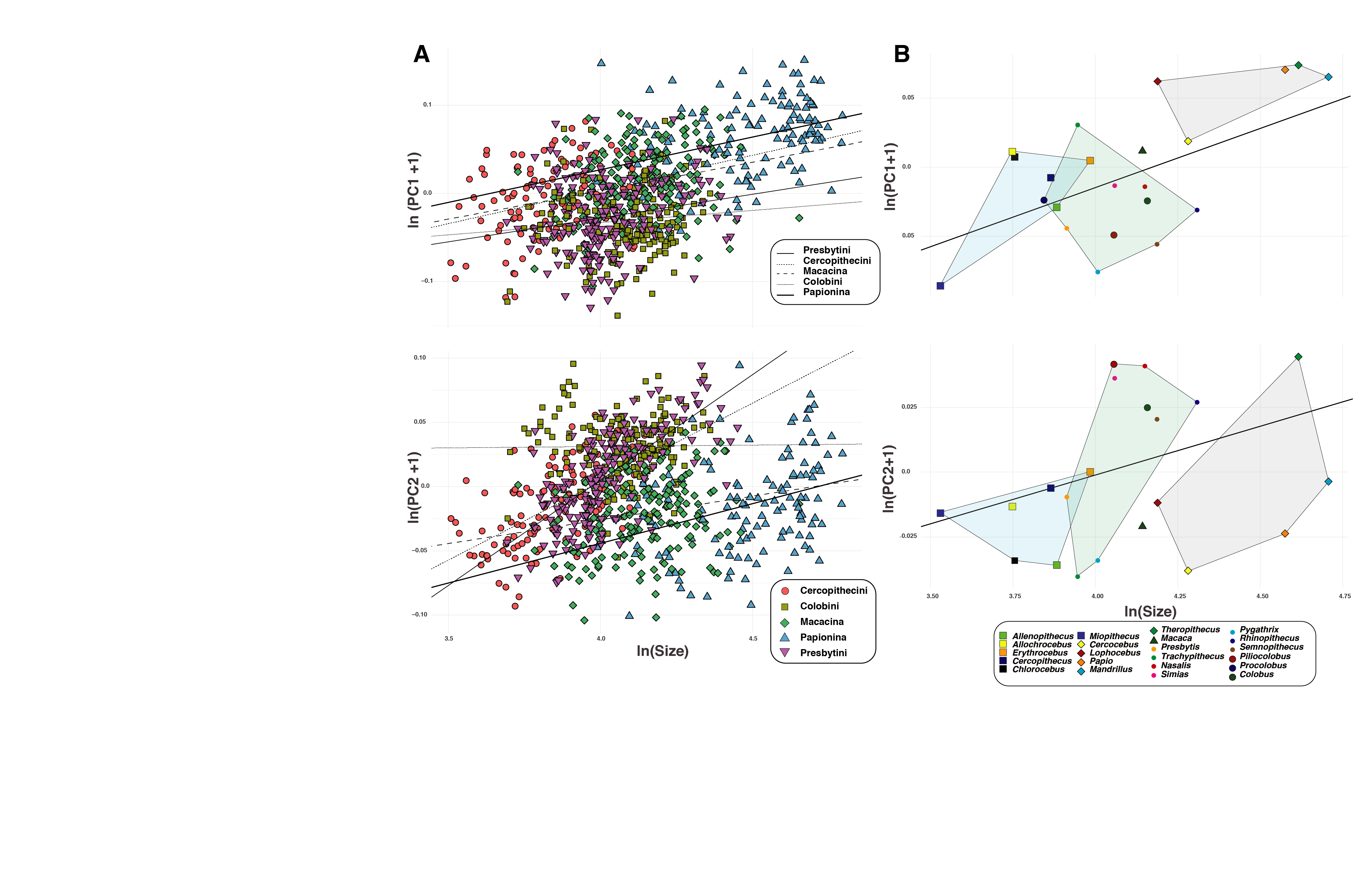

### Figure_6.png

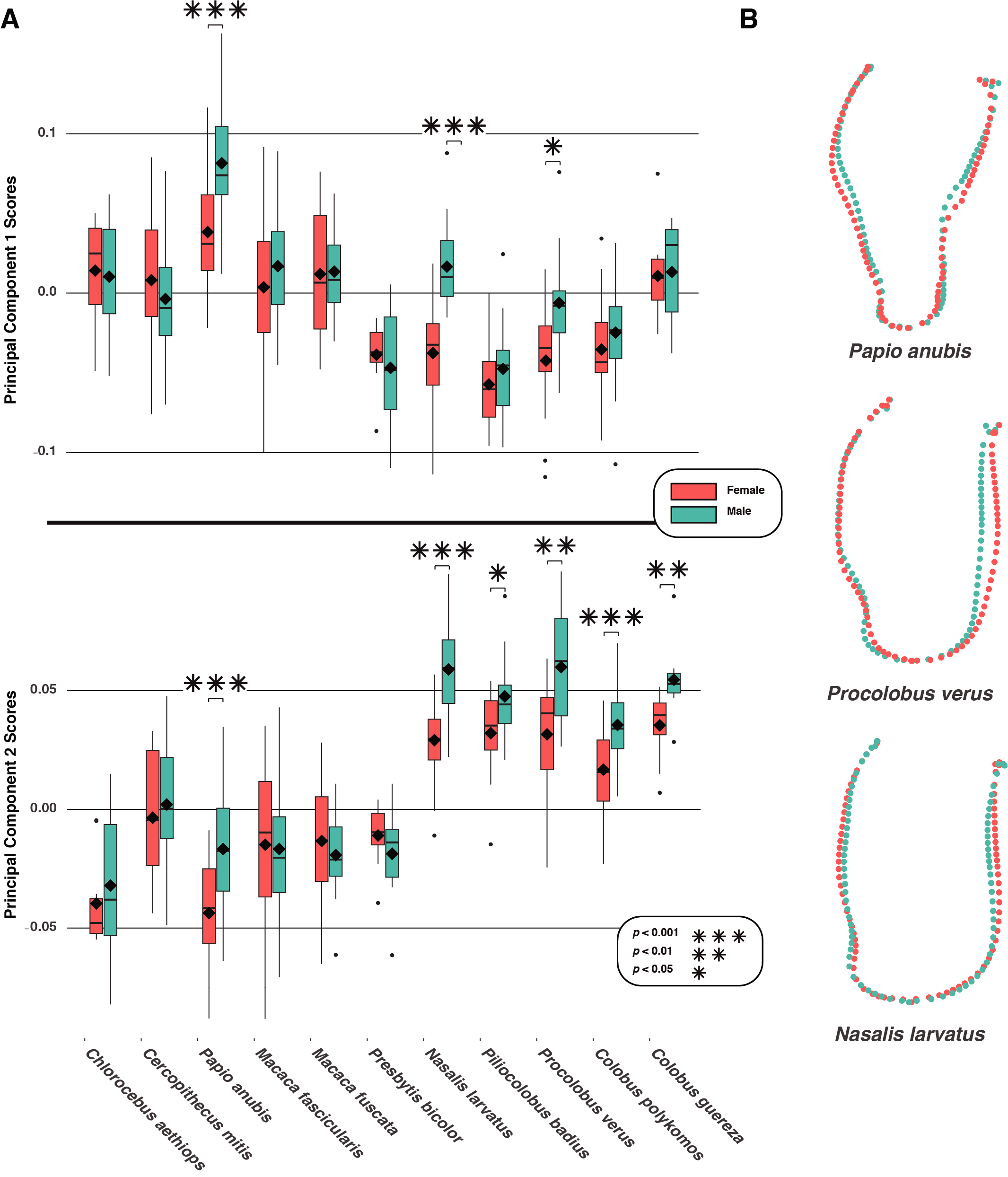
